# Investigating the NRAS 5’ UTR as a Target for Small Molecules

**DOI:** 10.1101/2022.01.05.475055

**Authors:** Sumirtha Balaratnam, Zachary R. Torrey, David R. Calabrese, Michael T. Banco, Kamyar Yazdani, Xiao Liang, Adrian R. Ferré-D’Amaré, Danny Incarnato, John S. Schneekloth

## Abstract

Neuroblastoma RAS (NRAS) is an oncogene that is deregulated and highly mutated in cancers including melanomas and acute myeloid leukemias. Constitutively activated NRAS induces the MAPK and AKT signaling pathways and leads to uncontrolled proliferation and cell growth, making it an attractive target for small molecule inhibition. Like all RAS-family proteins, it has proven difficult to identify small molecules that directly inhibit the protein. An alternative approach would involve targeting the NRAS mRNA. The 5′ untranslated region (5′ UTR) of the NRAS mRNA is reported to contain a G-quadruplex (G4) that regulates translation of *NRAS* mRNA. Stabilizing the G4 structure with small molecules could reduce NRAS protein expression in cancer cells by impacting translation. Here we report a novel class of small molecule that binds to the G4 structure located in the 5′ UTR of the NRAS mRNA. We used a small molecule microarray (SMM) screen to identify molecules that selectively bind to the NRAS-G4. Biophysical studies demonstrated that compound **18** binds reversibly to the NRAS-G4 structure with submicromolar affinity. A Luciferase based reporter assay indicated that **18** inhibits the translation of NRAS via stabilizing the NRAS-G4 *in vitro* but showed only moderate effects on the NRAS levels *in cellulo*. Rapid Amplification of cDNA Ends (RACE), RT-PCR analysis on 14 different *NRAS*-expressing cell lines, coupled with analysis of publicly available CAGE seq experiments, revealed that predominant NRAS transcript does not possess the G4 structure. Further analysis of published rG4 and G4 sequencing data indicated the presence of G4 structure in the promoter region of *NRAS* gene (DNA) but not in the mRNA. Thus, although many *NRAS* transcripts lack a G4 in many cell lines the broader concept of targeting folded regions within 5’ UTRs to control translation remains a highly attractive strategy and this work represents an intriguing example of transcript heterogeneity impacting targetability.

## Introduction

Messenger RNA (mRNA) transcripts exhibit highly complex and diverse structures through canonical and non-canonical base pairing interactions as well as interactions with proteins^1-4^. Structured RNA elements impact biological functions such as RNA synthesis, metabolism, and regulatory pathways^1, 5-6^. The formation of folded structures in mRNA has been shown to play important roles in post-transcriptional regulation of gene expression including RNA maturation, translation, and degradation^7-9^. Interestingly, folds in 5′ untranslated regions (UTRs) of mRNAs have been recognized as a major feature that regulates the translation process^10-11^. For example, ∼60% of 5′ UTRs in humans have structured regions near the 5′ cap site and are believed to impact translation initiation^12^. One way these structures regulate cap-dependent translation initiation through helicase-mediated remodeling of RNA structures and higher-order RNA interactions, as well as cap-independent translation initiation through internal ribosome entry sites (IRESs)^10-11, 13^. The formation of complex structures within the 5′ UTRs suggests opportunities for small molecule targeting and control of gene expression at the translational level.

NRAS is a proto-oncogene belonging to the RAS oncogene superfamily and was first identified in neuroblastoma^14-16^. In many cancers, NRAS proteins are constitutively activated by oncogenic mutations or overexpression. The mutation of NRAS accounts for about 15% of RAS related human malignancies, notably in myeloid leukemias and cutaneous melanomas^17-19^. In normal cells, NRAS proteins switch between active GTP-bound forms and inactive GDP-bound forms. The transition between the active and inactive state is mediated by GTPase-activating proteins^20-21^. Oncogenic NRAS hyperactivation leads the persistence of the GTP-bound state of NRAS and initiates constitutive MAPK signaling as well as AKT signaling, driving malignancies by promoting cell growth, survival, and invasion^22-25^. Targeting NRAS is therefore believed to be a promising strategy for developing anticancer therapies. However, the NRAS protein has been considered a highly challenging target due to a lack of drug binding pockets other than nucleotide (GTP) binding pocket^19, 25-26^. Since the nucleotide binding pocket has picomolar binding affinity to GTP, developing GTP-competitive inhibitors is nontrivial. Still, the FDA approval of the covalent KRAS^G12C^ inhibitor AMG 510 validates Ras-family proteins as highly important anticancer targets^27^ even though there are no approved NRAS inhibitors yet. One intriguing strategy to expand the targetability of NRAS is to target the associated mRNA and inhibit translation. This approach could be accomplished by identifying a structured region that contain small-molecule binding pockets in the NRAS mRNA and developing the small molecules that bind to these structures.

G-quadruplexes (G4s) are a class of secondary structure formed by G-rich sequences in DNA or RNA^28^. The general formula for canonical^29^ G4 forming sequences is G _2-5_ N _1-7_ G _2-5_ N _1-7_ G _2-5_ N _1-7_ G _2-5_ where N can be any nucleotide located in the loops of the G4. One G from each of the four G-tracts is bonded to form a co-planar G-quartet by Hoogsteen base pairings. Two or more G-quartets can stack on top of each other to form G4 structures. G4s are stabilized by monovalent cations, most commonly K^+30^. Although initial genome-wide studies revealed the existence of DNA-G4 structures in genomes and their enrichments in telomeric regions, the occurrence of G4s in RNA has been more controversial^31^. Studies have shown the presence of RNA-G4 structures in mRNAs, viral genomic RNA, pre-miRNAs, precursor piRNA transcripts, mature piRNAs and long noncoding RNAs (lncRNAs)^32^, while other reports also indicate that rG4s are globally unfolded in the steady state (presumably by helicases or other protein factors)^33-35^.

Still, rG4s are thought to play important roles in diverse of biological processes and related to human diseases^36-37^. One bioinformatic search for “regular” rG4s in 5′ UTRs of the human transcriptome indicated that 9,979 5′ UTRs contain at least one potential G4 forming sequence^38^. Further studies reported that formation of RNA-G4 structures in 5′ UTRs inhibit mRNA translation by interfering with the recruitment of the pre-initiation complex^39^. In the human *NRAS* gene, Kumari and co-workers reported the presence of RNA-G4 structure in the 5′ UTRs and that this G4 motif is conserved in both its sequence and its positions relative to the translation start site among different organisms such as chimpanzee, macaque, mouse, rat, and dog^40^. They used a luciferase reporter assay to demonstrate that formation of RNA-G4 structure in the 5′ UTRs of NRAS mRNA inhibits translation by >80% in rabbit reticulocyte lysates. This result suggested that native RNA-G4 structures in 5′ UTRs could act as regulatory elements of translation and become an attractive secondary structure for therapeutic targeting to control NRAS mRNA at the post-transcriptional level. Thus, stabilization of the NRAS-G4 represents an opportunity for pharmacological suppression of NRAS expression with small molecule ligands. The development of a few small molecule stabilizers for NRAS-G4 has been reported in recent years, though selectivity remains a challenge^41-42^.

In this study, we report the discovery of a series of small molecules that bind directly to the NRAS-G4 structure using an SMM screening strategy. The lead compound **18** showed selective binding to the NRAS-G4 structure over other RNA structures. Biophysical and biochemical experiments confirmed reversible submicromolar binding affinity of the lead compound. Additionally, luciferase-based reporter assays indicated that compound inhibits translation in rabbit reticulocyte lysates via stabilizing the NRAS-G4 structure *in vitro*. Structure probing and X-Ray crystallographic studies support the formation of a parallel-stranded G4 in the context of the luciferase reporter construct. Treatment of **18** in SK-MEL-2 and MCF-7 cells resulted in a moderate effect on NRAS translation (∼ 20%). In-depth analysis of *NRAS* transcripts in various cell lines using RT-PCR and 5′ RACE confirmed that majority of NRAS transcript isoforms are shorter and lack a G4-forming sequence in the 5′-UTR. This result was further supported by analysis of publicly available rG4 sequencing and CAGE seq data and aligns with the recently re-annotated *NRAS* transcription start site in the UCSC genome database^43-47^. Importantly, this work does not rule out the existence of an NRAS rG4, but rather demonstrates that the predominant NRAS transcript lacks it in all the cell lines evaluated and represents an intriguing example of how heterogeneity in transcript length can impact the ability to target an mRNA. Finally, this work demonstrates that the strategy of controlling translation by targeting structured regions of 5’ UTRs is a viable approach.

## Results

### Discovery of an NRAS G4-binding Small Molecule using SMM

To identify small molecules that bind to the NRAS-G4, we used an SMM screening strategy^48^. SMM is a convenient method to rapidly screen tens of thousands of compounds to identify selective RNA-binding small molecules and has successfully been employed by our lab and others to identify RNA-binding ligands^48^. In the iteration of this approach used by our laboratory, a library of 26,227 compounds is spatially arrayed and covalently linked to the glass surface. An AlexaFluor647-labeled NRAS-G4 oligonucleotide was designed as a screening construct, folded into a G4 structure in K^+^ buffer, and incubated with printed arrays (Figure 1A). The fluorescence intensity is measured at each location on the array to identify binding interactions between the NRAS-G4 and the compound printed at each location. A composite Z-score is then calculated for each compound (printed in duplicate) in the library. Next, Z-scores for NRAS-G4 the incubated data set are compared to a control (buffer incubated) data set. Compounds are considered hits if (1) they had a composite Z-score greater than three (2) Coefficient of Variation (CV) of the two compound spots was lower than 200% and (3) compounds showed an increase in fluorescence in the presence of the RNA compared to the buffer incubated slides^48^. We identified 235 hits from the compounds screened, for an overall hit rate of 0.89%. To further investigate selectivity, Z-scores for each hit compound were compared to 20 different SMM screens that were performed with various RNA and DNA structures, including 11 other DNA or RNA G4s (Supplementary information Table S1). After selectivity analysis, 30 unique hit compounds were identified as having high Z score and good selectivity (Supplementary information Table S2). As a representative example, direct binding on SMM slides (Figure 1B) and selectivity data (Figure 1C) are shown for compound **1**. From the set of NRAS-G4 selective hit molecules, 14 compounds were selected for further analysis based on their availability and chemotype (supplementary information Table S3 and Figure S1). To validate the 14 hit compounds as NRAS-G4 binders, binding assays were performed by Surface Plasmon Resonance (SPR) (Supplementary information Table S3). Based on the measured equilibrium constant (K_D_), compound **1** was chosen for further analysis.

**Figure 1:**
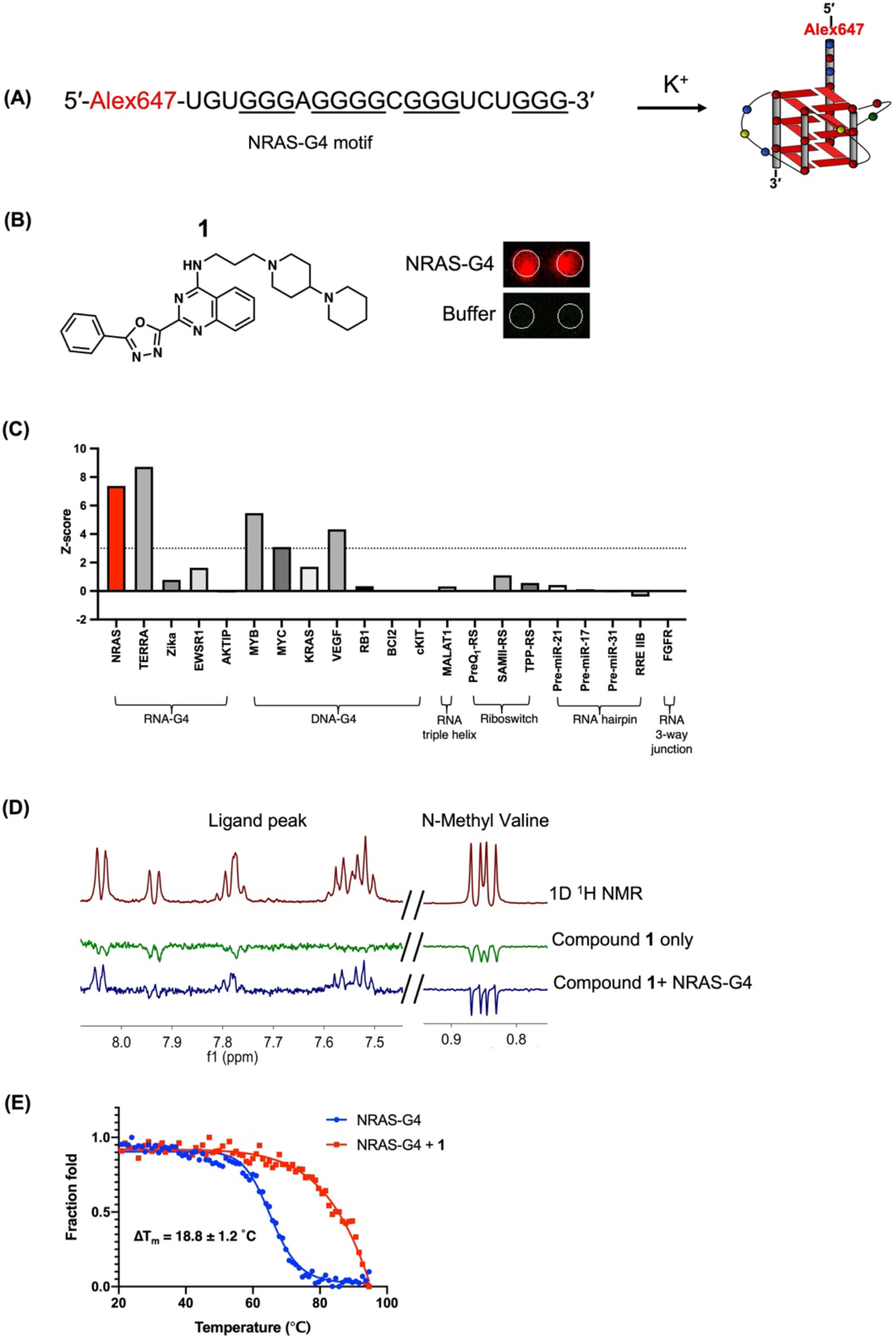
Small Molecule Microarray (SMM) screening using the NRAS-G4 and hit validation. **(A)** Sequence of the AlexaFluor647-labeled NRAS G-quadruplex forming sequence (5′-Alex647-NRAS-G4) used for SMM screening. Schematic drawing of parallel G4 structure formed by the folded sequence in the presence of K^+^ ion. **(B)** SMM images and chemical structure of hit compound **1**. Note that compounds are printed in duplicate. **(C)** Selectivity of the hit compound **1** across 20 different oligonucleotides screened using the SMM platform as measured by composite Z-score. **(D)** NMR validation of hit compound **1**. The ^1^H NMR of **1** and N-methyl-L-valine (non-binding control) (Top, red spectrum), WaterLOGSY NMR of **1** and N-methyl-L-valine in the absence (middle, green spectrum) and presence (bottom, blue spectrum) of NRAS-G4 RNA. **(E)** Representative thermal melting curve of NRAS-G4 in the absence (blue) and presence (red) of compound **1**. Melting was performed using 1 mM K^+^. Error bars indicate the standard deviation determined from three independent measurements.

### Biophysical analysis confirmed the binding of compound 1 to NRAS-G4

We first evaluated the binding interaction between the initial hit compound **1** and the NRAS-G4 using ligand-observed NMR experiments. To determine the solubility of the compound in the aqueous buffer, compound **1** was observed in a standard ^1^H NMR experiment (Figure 1D upper panel). Next, WaterLOGSY was performed on **1** both in the presence and absence of the NRAS-G4. As shown in the Figure 1D, in the absence of RNA, all peaks phased negatively, confirming the compound does not aggregate in aqueous buffer. Upon the addition of NRAS-G4 RNA, peaks corresponding to **1** phased positively, while peaks for N-methyl-L-valine (used as an internal, non-binding control) remain unchanged in all cases confirming the direct binding of **1** to the NRAS-G4.

To measure the binding affinity of compound **1** with NRAS-G4, we used surface plasmon resonance (SPR) experiments. A biotinylated NRAS-G4 oligonucleotide was immobilized to a streptavidin-coated SPR chip, and binding was measured as a function of compound concentration. This experiment demonstrated that compound **1** bound to NRAS-G4 with an equilibrium dissociation constant (K_D_) of 0.45 ± 0.11 μM (Supplementary information Figure S2 A and B). To validate binding further using an orthogonal biophysical assay, a fluorescence titration assay was also performed to measure the affinity of **1** to NRAS-G4. In this experiment, compound **1** was titrated into a solution containing AlexaFluor 647-labelled NRAS-G4 (the same construct used in the SMM screen) and changes in fluorescence intensity were monitored as a function of compound concentration (Supplementary information Figure S2 C). An apparent dissociation constant was measured by fitting the binding curve between the normalized fluorescence intensity and compound concentration. In this assay, compound **1** had a K_D_ of 1.2 ± 0.5 μM. In sum, biophysical experiments confirmed the direct binding of **1** to the NRAS-G4 structure and good solubility in aqueous buffer.

Next, we evaluated effect of **1** on NRAS-G4 thermal stability by performing a circular dichroism (CD)-based thermal melt assay, in which molar ellipticity was measured as a function of increasing temperature. In the absence of **1**, the CD spectrum of the NRAS-G4 exhibited a maximum at 263 nm and a minimum at 240 nm, confirming proper folding of a parallel G4 structure^40^. The melting temperature (T_M_) of the RNA in 1 mM K^+^ was measured to be 65.4 ± 1.9 °C, which was consistent with previously reported values^40^. In the presence of 2.5 µM of compound **1**, the T_M_ was also increased by 6.9 ± 0.6°C, while at 10 μM of compound **1** increased the T_M_ by 18.8 ± 1.2 °C (Figure 1E and supplementary Figure S3).

### Structure-activity relationship (SAR) study of 1

To identify an improved binder, we performed a preliminary structure-activity relationship (SAR) study by using a focused series of commercially available analogs of **1** (Table 1). Based on the chemical structure of compound **1**, 13 analogs of compound **1** were purchased from the commercially available focused library. These analogs have altered R^1^ side chain groups, while R^2^ and R^3^ were also altered in several analogs. Each compound was evaluated for binding affinity towards the NRAS-G4 structure by fluorescence titration assay (Figure 2A and supporting information Figure S4). Most of the analogs showed strong binding behavior toward the NRAS-G4 structure and 12 analogs out of 13 (except compound **8**) were identified as stronger binders than the parent compound (**1**). Among them compounds **2, 14** and **18** showed the tightest binding affinities and the calculated binding affinity was less than 300 nM. To further validate the binding affinities of these compounds, SPR experiments was performed (Figure 2B, C and supporting information Figure S5). Here, analogs **2** and **14** showed weaker binding affinity with 4.5 ± 1.2 μM and > 41 μM respectively. However, compound **18** bound to NRAS-G4 with an equilibrium dissociation constant (K_D_) of 932 ± 38 nM in SPR compared to a 250 nM K_D_ in fluorescence titrations. Based on these results, we selected **18** as a lead compound for further analysis. To examine whether compound **18** influenced the stability to the NRAS-G4 structure, we performed a CD-based thermal melting assay. In the presence of compound **18**, the melting temperature of NRAS-G4 increased by 6.2 ± 0.4 °C (Figure 2D).

**Table 1:** Focused library of analogs of compound **1** and affinity for the NRAS G4

| Name | R<sup>1</sup> | R<sup>2</sup> | R<sup>3</sup> | FIA K<sub>D</sub><br>(μM) | Name | R<sup>1</sup> | R<sup>2</sup> | R<sup>3</sup> | FIA K<sub>D</sub><br>(μM) |
| --- | --- | --- | --- | --- | --- | --- | --- | --- | --- |
| 1 |  | -H | -H | 1.2 ± 0.5 | 12 |  | -H | -CH<sub>3</sub> | 0.30 ± 0.06 |
| 2 |  | -F | -H | 0.26 ± 0.07 | 14 |  | -H | -CH<sub>3</sub> | 0.28 ± 0.04 |
| 3 |  | -F | -H | 0.43 ± 0.12 | 16 |  | -CH<sub>3</sub> | -H | 0.35 ± 0.05 |
| 6 |  | -H | -H | 0.74 ± 0.03 | 17 |  | -CH<sub>3</sub> | -H | 0.56 ± 0.17 |
| 7 |  | -H | -H | 0.36 ± 0.06 | 18 |  | -CH<sub>3</sub> | -H | 0.25 ± 0.06 |
| 8 |  | -Cl | -H | 3.0 ± 1.7 | 24 |  | -CH<sub>3</sub> | -H | 0.50 ± 0.14 |
| 10 |  | -H | -CH<sub>3</sub> | 0.48 ± 0.08 | 25 |  | -CH<sub>3</sub> | -H | 0.95 ± 0.45 |

**Figure 2:**
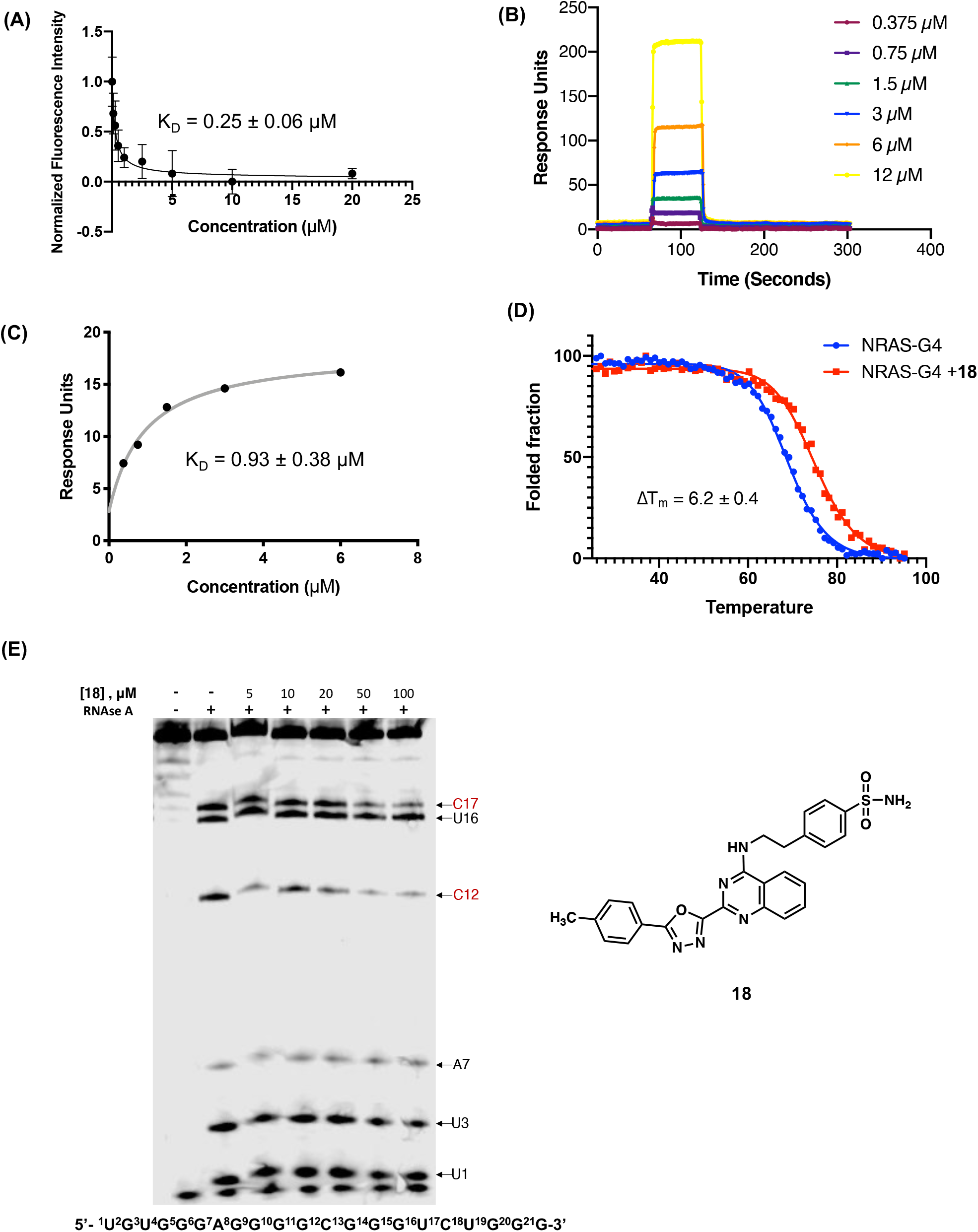
Affinity of compound **18** to the NRAS-G4. **(A)** Fluorescence intensity assay of 5′-(AlexaFluor647)-labeled NRAS-G4 RNA in the presence of **18**. Error bars indicate the standard deviation determined from three independent measurements. **(B)** Sensorgrams and **(C)** binding curve corresponding to **18** interacting with a 5′ biotin labeled NRAS-G4 RNA in SPR. **(D)** CD-based thermal melting of NRAS-G4 in the absence (blue) and presence (red) of 10 μM compound **18** in 1 mM K^+^ buffer. **(E)** RNase A structure probing of NRAS-G4 with increasing concentration of compound **18**. Compound **18** protected nucleotides (C17 and C12) in the loops (labelled in red). Chemical structure of compound **18**.

The binding selectivity of compound **18** to NRAS-G4 over other G4 structures was also evaluated by FIA. For this study, nine different G4 structures from cancer-relevant genes were selected. This panel includes 6 DNA G4s from the promoter region of oncogenes (*BCL2, KRAS, mTOR*, Telomeric DNA G4, *VEGF*, and *MYCN*) and 3 RNA-G4s from mRNA 5′ UTRs or ncRNAs (AKTIP, TERRA and EWSR1) (Supplementary information Table S4). Compound **18** was titrated into the various 5′-Cy5 or AlexaFluor647 labeled DNA/RNA G4s and apparent K_D_ values were calculated. We observed that compound **18** exhibited weak binding towards the RNA-G4 structure formed by AKTIP and DNA-G4 from BCL2, (Supplementary information Figure S6). For all other G4s, the binding affinity (K_D_) could not be calculated because **18** showed no significant quenching. These results, combined with selectivity observed in SMM assays (Figure 1C) indicate that **18** binds to the NRAS-G4 via its unique structure.

### Enzymatic structure mapping indicates a binding site of 18 on NRAS-G4

To biochemically investigate the binding mode of the compound **18** on the NRAS-G4 structure, we performed Ribonuclease A (RNAase A) structure mapping^49^. A 5′-(AlexaFluor647)-labeled NRAS-G4 RNA (5 μM) was folded into a G4 structure and incubated with increasing concentrations of compound **18** and RNase A. As shown in Figure 2E, in the absence of **18**, RNAse A predominantly cleaved the U and C bases in the loop regions of G4 structure. Upon incubation with increasing concentration of **18** however, the resulting RNAse A cleavage pattern was markedly different at C12 and C17, flexible bases anticipated to be on neighboring loops (Figure 2E and Supplementary Figure S7). The C12 nucleotide was protected from RNAse A cleavage at lower concentrations of **18** (5μM), while C17 protection was only observed at higher concentrations of **18** (50μM and 100μM). These effects indicate that compound **18** binding impacts the flexibility or accessibility of the loop regions and presumably indicate the binding site of **18** on the NRAS-G4 structure.

### Compound 18 inhibits the translation of an NRAS reporter gene *in vitro*

To evaluate the effect of compound 18 on the efficiency of NRAS translation, we utilized a reporter system developed by the Balasubramanian group.^40^ Briefly, we used a plasmid containing 254 bp encoding the human NRAS 5′ UTR mRNA upstream of a T7 promoter (pSKC11), which possesses the transcript containing the G4 structure (NRAS-G4-FL) (Figure 3A). As a control, another plasmid was used lacking the first 29 bp of the 5′ UTR to produce a transcript that lacks the G4-forming region (NRAS-G4-Del-FL). Each of the transcripts was generated by *in vitro* transcription using T7 RNA polymerase, and the transcripts were subjected to *in vitro* translation in rabbit reticulocyte lysates in the presence of compound **18**. The efficiency of translation was measured by quantifying the resulting photon output and normalizing to DMSO controls. As shown in Figure 3B, dose-dependent translational inhibition was observed with NRAS-G4-FL in the presence of compound **18**. However, no translational inhibition was observed for **18** with the control NRAS-G4-Del-FL transcript. Thus, compound **18** only inhibits the translation of the reporter gene containing the NRAS 5’UTR when the G4 is present.

**Figure 3:**
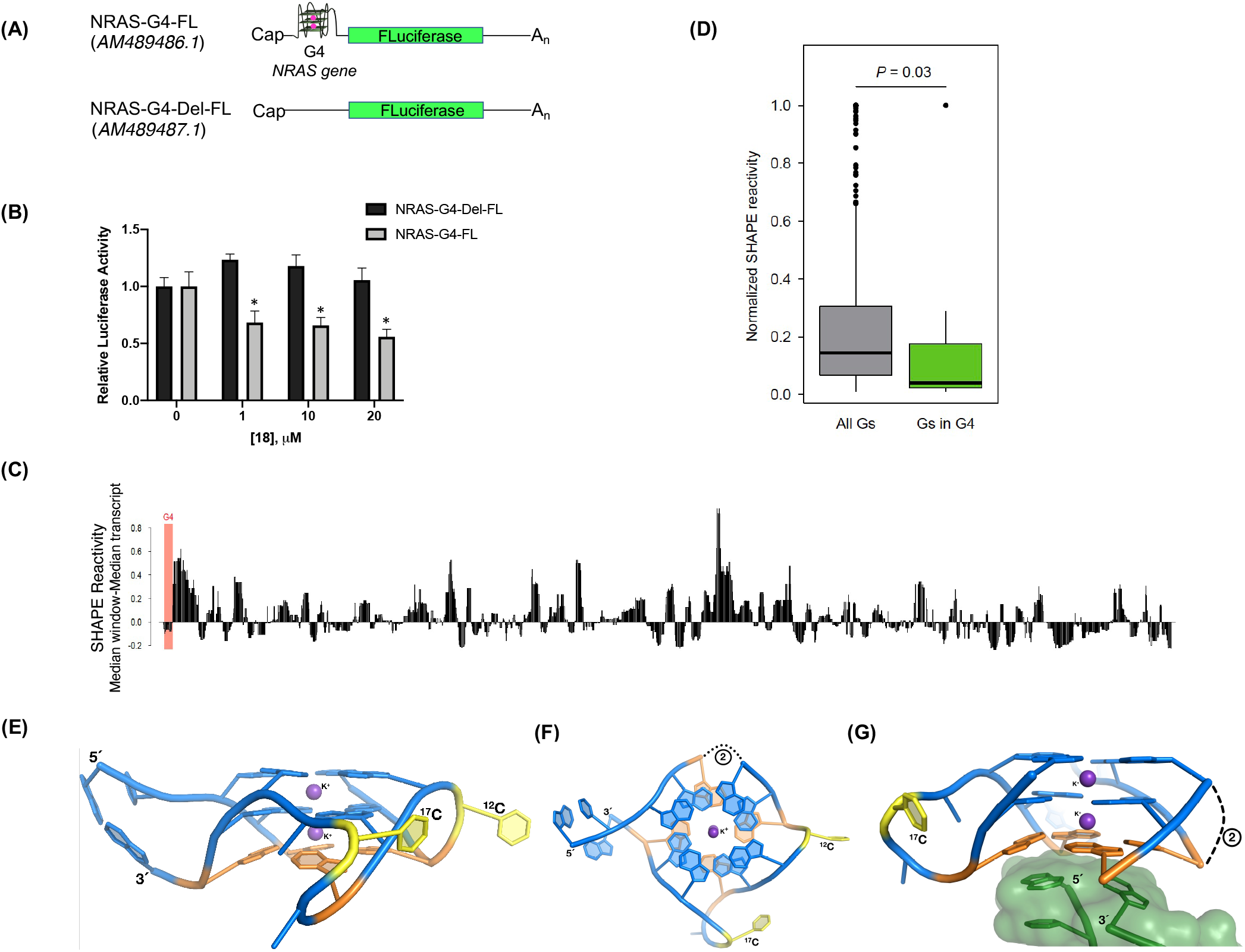
Effect of **18** on NRAS-FL reporter gene translation and structural analysis of the NRAS G4. **(A)** Schematic representation of firefly luciferase reporter constructs that contain the NRAS 5′ UTR: NRAS-G4-FL (contains G4, top) and NRAS-G4-Del-FL(lacks G4, bottom). **(B)** Relative translation efficiency of constructs in the presence of increasing concentrations of **18**, measured by quantitation of luciferase enzyme activity. Results were normalized to data for constructs with DMSO. Error bars represent the standard deviation of three independent experiments (n=3, *P < 0.05). **(C)** Median SHAPE reactivities across the chimeric luciferase construct (NRAS-G4-FL). Regions below the X axis indicate more structure than average, while regions above the X axis indicate less structure than average. **(D)** Bar graph represents normalized SHAPE reactivity of all Gs and Gs involved in the G4 formation. **(E-F)** Cartoon representation of the X-ray crystal structure of the NRAS G4 viewed on the G tetrads plane **(E)** and rotated 90° **(F)**. The RNA is colored blue, except the buckled 3’ quartet (orange) and loop nucleotides (yellow).

To confirm formation of the NRAS-G4 structure is formed in the context of the chimeric reporter construct and the observed *in vitro* translation effect was due to the G4 binding of **18**, we performed the Selective 2′-Hydroxyl Acylation Analyzed by Primer Extension-Mutational Profiling (SHAPE-MaP) experiments^50^. SHAPE-MaP is an RNA structure probing technique that combines chemical probing of unpaired nucleotides with next-generation sequencing to measure the flexibility of individual nucleotides in long RNAs. We treated pre-folded NRAS-G4-FL mRNA construct with the SHAPE reagent NAI, and SHAPE-MaP was performed to generate a reactivity profile for the entire chimeric luciferase reporter construct (NRAS-G4-FL) (Supplementary information S8). SHAPE reactivity indicates the relative flexibility of a nucleotide, which correlates with occurrence of base-pairing and base stacking (Supplementary information Figure S8). SHAPE reactivities above 0.8 indicate likely unpaired bases, while reactivities below 0.4 indicate likely base-paired nucleotides. As shown in figure 3C, the median window SHAPE reactivity was plotted across the transcript using a sliding window (median window-median transcript). This analysis demonstrated that there are specific regions in the transcript that have low median SHAPE reactivity. Most importantly, the SHAPE reactivities of the G4 forming region (highlighted in red) revealed that this region was structurally constrained and are involved in a folded structure.

Comparison of the normalized SHAPE reactivities of the guanine nucleotides involved in the formation of G4 structure with all other guanines nucleotides in the chimeric luciferase construct demonstrated the guanines involved in G4 formation have low SHAPE reactivity (Figure 3D). The observed low SHAPE reactivity values indicate a high degree of ordered structure in the transcript, presumably corresponding to a folded G4. Interestingly, G12 showed higher SHAPE reactivity compared to the other guanines in the NRAS-G4 forming sequence. Thus, the dominant NRAS-G4 structure is likely to consist of G12 located in a loop region connecting the G-quartets and not involved in the formation of the G-quartet. Efforts to map compound binding to this mRNA did not meet with success, potentially because the G4 is already highly folded in the absence of compound.

In order to gain molecular insights into the structure of the G4, we determined the X-ray crystal structure of a NRAS-G4 variant at 2.9 Å resolution (Figure 3E,F and supplementary Table S5). To prevent conformational heterogeneity of NRAS-G4 structural isomers, residue G8 was mutated to ensure formation of a single conformer. A single molecule of NRAS-G8U was observed in the asymmetric unit. The NRAS-G8U structure adopts a canonical parallel quadruplex that is comprised of three G-quartets coordinated by potassium ions. The guanines of the G4 are in the anti-conformation but consist of a mixture C2’- and C3’-endo sugar puckers (Supplemental Figure S9). In the NRAS-G8U structure, the G-tracts are connected by propeller-type loops. The loop nucleotides corresponding to A7 and U8 lacked electron density, were presumed to be disordered, and thus were not modeled. The other loops of the G4 possesses electron density, which includes C12 and C17. In the crystal structure, the 3’ G-quartet exhibits substantial buckling, whereas the other two G-quartets form largely planar tetrads (Figure 3G). The buckled guanosines of the 3’ G-quartet adopt a C2’-endo pucker, which deviates from the preferred C3’-endo pucker observed in most RNA G4s^29^. This bulked quartet participates in crystal packing; thus its conformation in solution may differ.

Potassium ions are depicted as purple spheres. Black dashes denote two disordered nucleotides not observed in the structure. **(G)** Interface between a NRAS-G8U molecule and an adjacent crystallographic asymmetric unit (ASU). A portion of the molecule from an adjacent ASU is highlighted by the translucent green molecular surface. The crystal contacts may contribute to bulking of the 3’ G-quartet (colored orange).

### Compound 18 has moderate effects on NRAS levels *in cellulo*

To examine the effect of compound **18** on cellular NRAS levels, we selected a breast cancer cell line (MCF-7) and a melanoma cell line (SK-MEL-2). Both cancer cell lines have high level of NRAS, either by oncogenic mutation (SK-MEL-2) or overexpression (MCF-7). Compound **18** was evaluated for its capacity to reduce cell viability in both cell lines. We observed that **18** decreased MCF-7 and SK-MEL-2 cell viability with an IC_50_ of 1.1 ± 0.3 μM and 0.9 ± 0.4 μM respectively (Figure 4A). To examine the effect of compound **18** on NRAS mRNA stability, we performed quantitative PCR assay (qPCR) to quantify the NRAS mRNA level after treatment of compound **18**. Treatment with **18** did not drastically change the NRAS mRNA levels in either MCF-7 or SK-MEL-2 cells up to 25 μM (Figure 4B). To assess the changes more broadly in the transcriptome with treatment of compound **18**, we performed gene expression profiling by RNA-seq. Both cell lines were treated with 1μM of compound **18** and total RNA was isolated from cells after 48 hours. RNA sequencing was performed, and differential gene expression analysis was carried out to compare DMSO and **18**-treated sample groups. This analysis identified 384 differentially expressed genes (290 downregulated and 94 upregulated genes) in MCF-7 cells and 292 genes (226 downregulated and 66 upregulated genes) in SK-MEL-2 cells with a false discovery rate (FDR)<0.05 (Figure 4C-D). As with qPCR analysis, compound **18** treatment did not change levels of NRAS mRNA in either cell line (Figure 4C-D and Supplementary information S10). We also observed that **18** had no effect on the levels of other RAS family genes (*KRAS* and *HRAS*). Additionally, ontology analysis (Supplementary Table S7 and S8) did not indicate perturbations to RAS-related signaling pathways in either cell line. To further evaluate the effect of compound **18** on other mRNAs which contain G4 structures in their UTR regions (3′-UTR and 5′-UTR) or CDS, we analyzed the FPKM (Fragments Per Kilobase of transcript per Million mapped reads) values in 29 different G4 containing mRNAs, including 24 mRNAs with a G4 in 5′-UTR, 3 with a G4 in the 3′-UTR and 2 containing G4s in the coding sequence. No significant change in expression was observed with compound **18** treatment in mRNAs which contain G4 structures either in their UTR regions (3′-UTR and 5′-UTR) or CDS (Supplementary information S11). Overall, the RNA seq analysis indicates that compound **18** does not broadly affect the expression of genes with G4 structures in mRNA or promoter (DNA).

**Figure 4:**
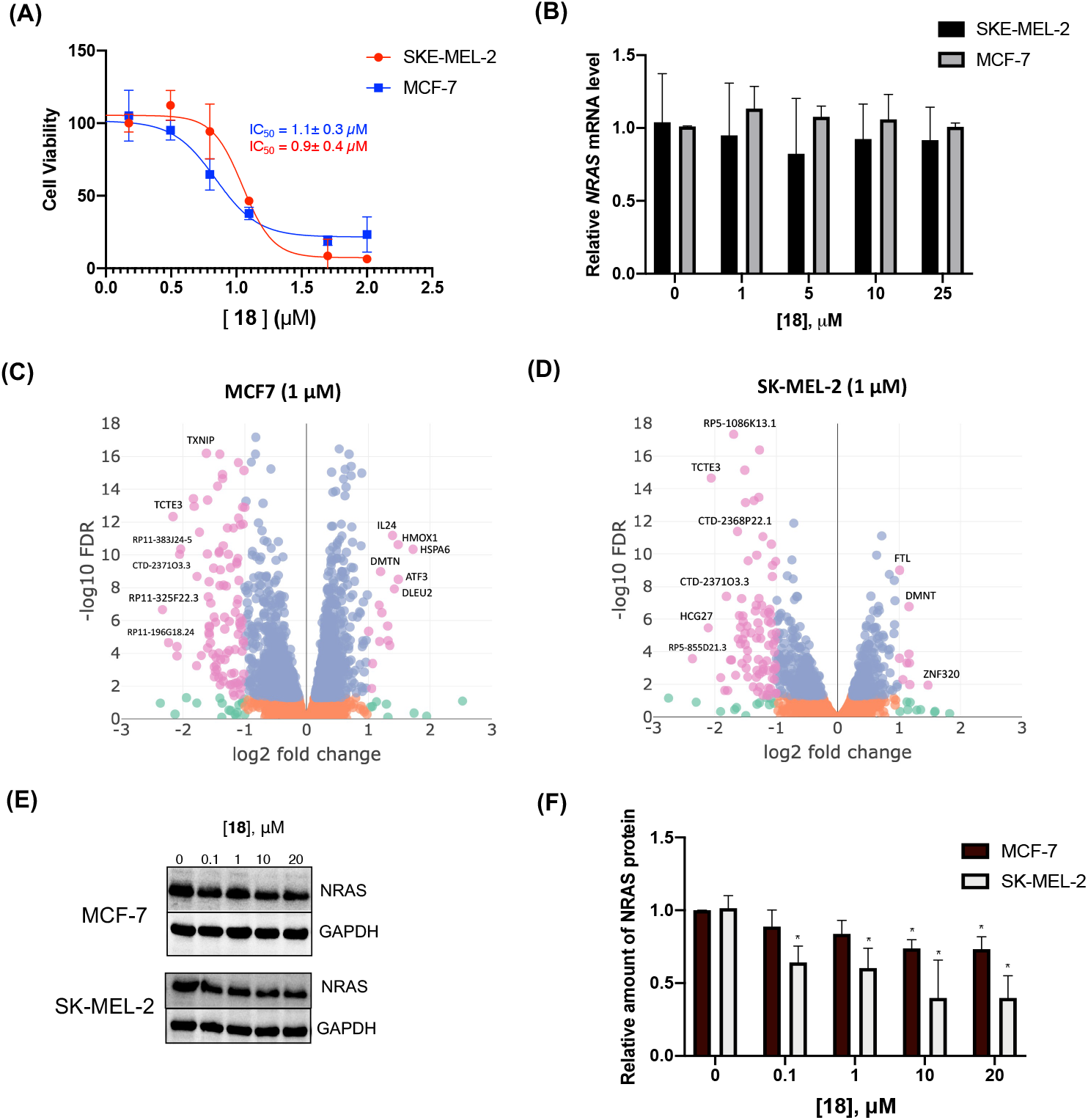
Effect of **18** on NRAS expression *in cellulo*. **(A)** Inhibition of SK-MEL-2 cell and MCF-7 cell proliferation with treatment of **18**. The results are presented as the mean ± SEM (n = 9) of three independent experiments with triplicate in each. **(B)** The histogram representing the relative level of *NRAS* mRNA level to *GAPDH* after **18** treatments in SK-MEL-2 and MCF-7 cells. The data were generated by qPCR analysis. The results are presented as the mean ± SEM (n = 3) of three independent experiments. **(C-D)** Volcano plot of differential gene expression analysis (DeSeq2) with 1μM compound **18** treatment in MCF-7 cells and SK-MEL-2 cells. Cells treated with DMSO act as a control. All analyses performed in 3 independent replicate samples. **(E)** Western blot of NRAS protein level after compound **18** treatment in SK-MEL-2 cells and MCF-7 cells. **(F)** Relative level of NRAS protein level to GAPDH in SK-MEL-2 cells and MCF-7 cells after **18** treatments quantified by densitometry analysis. The statistical significance was calculated by t-test analysis. (n=3, *P < 0.05).

Next, to investigate the functional inhibition of **18** on NRAS translation, MCF-7 and SK-MEL-2 cells were treated with increasing concentration of compound **18** (0.1 to 20 μM). As shown in Figure **4E-F**, 20 μM **18** caused a modest 20-30% decrease in NRAS levels at higher concentrations. Since we observed a weaker effect of compound **18** on NRAS levels in cells than in the luciferase reporter *in vitro*, we performed an in-depth analysis of 5′-UTR structure of the endogenous NRAS transcript.

### Analysis of the architecture of 5′-UTR of NRAS mRNA

Based on NCBI annotation, two different transcripts for the NRAS mRNA exist: an older version (NM_002524.4) and a revised, newer version (NM_002524.5). The original NRAS transcript annotation contained a 4454 nt long sequence including a 254 nt 5′-UTR which contains the G4 (between15-32 nt from the 5′-end). However, a revised annotation of the NRAS transcript is 4326 nt long and has a 131 nt long 5′-UTR region that lacks 123 nt from the 5′-end of the older transcript. Importantly, the revised annotation of the NRAS transcript is devoid of a G4 structure in its 5′-UTR. Transcripts with differences in 5′UTR length are not uncommon due to the presence of multiple promoters, alternative transcription start sites, alternative splicing mechanisms within UTRs, or inaccurate mapping^51^.

To experimentally define the NRAS mRNA 5′-UTR, we performed a series of RT-PCR experiments. We designed primer pairs A/B that specifically bind and amplify the G4-containing 5′-UTR of the NRAS mRNA and C/D pair that bind and amplify the part of the coding region of the NRAS mRNA (Figure 5A). The chimeric luciferase construct transcript (NRAS-G4-FL) acted as a positive control for this experiment as it contains the G4 in its 5′-UTR. For this control, E/F primers were also designed to amplify the coding region of the luciferase mRNA. As Shown in figure 5B-C, 14 different cell lines were analyzed, all of which contained measurable levels of *NRAS* mRNA. In the control system (NRAS-G4-FL), both the G4-containing and coding sequence primers were amplified equally well. However, all the cell lines examined showed only trace amplification of the G4 containing 5′-UTR (Figure 5B-C, indicated with red arrows), while in all cases the coding sequence primers were amplified with similar intensity to the control NRAS-G4-FL. These results indicate that the majority of NRAS transcripts are devoid of a G4 structure in the 5′ UTR. To further confirm this observation, we performed 5′ rapid amplification of cDNA ends (RACE) experiments to map the 5′ end of the NRAS mRNA in HEK-293 cells^52^. RACE is a powerful PCR-based technique for the rapid characterization of the 5′ end of mRNAs and the start of transcription. As shown in figure 5D, RACE experiments confirm that predominant NRAS transcript in HEK-293 cells lacks a G4 structure and the G4 forming regions are upstream from the transcription start site (TSS). To further validate these observations, we analyzed publicly available Cap Analysis of Gene Expression Sequencing (CAGE-Seq) datasets of the *NRAS* transcript from several cell lines. CAGE-Seq is used to accurately annotate the 5′ end of RNAs carrying a cap-site and utilizes a “cap-trapping” technology. CAGE-Seq data also confirm that the major *NRAS* transcript lacks a G4 structure and TSS are located downstream to the G4 (Supplementary information S12).

**Figure 5:**
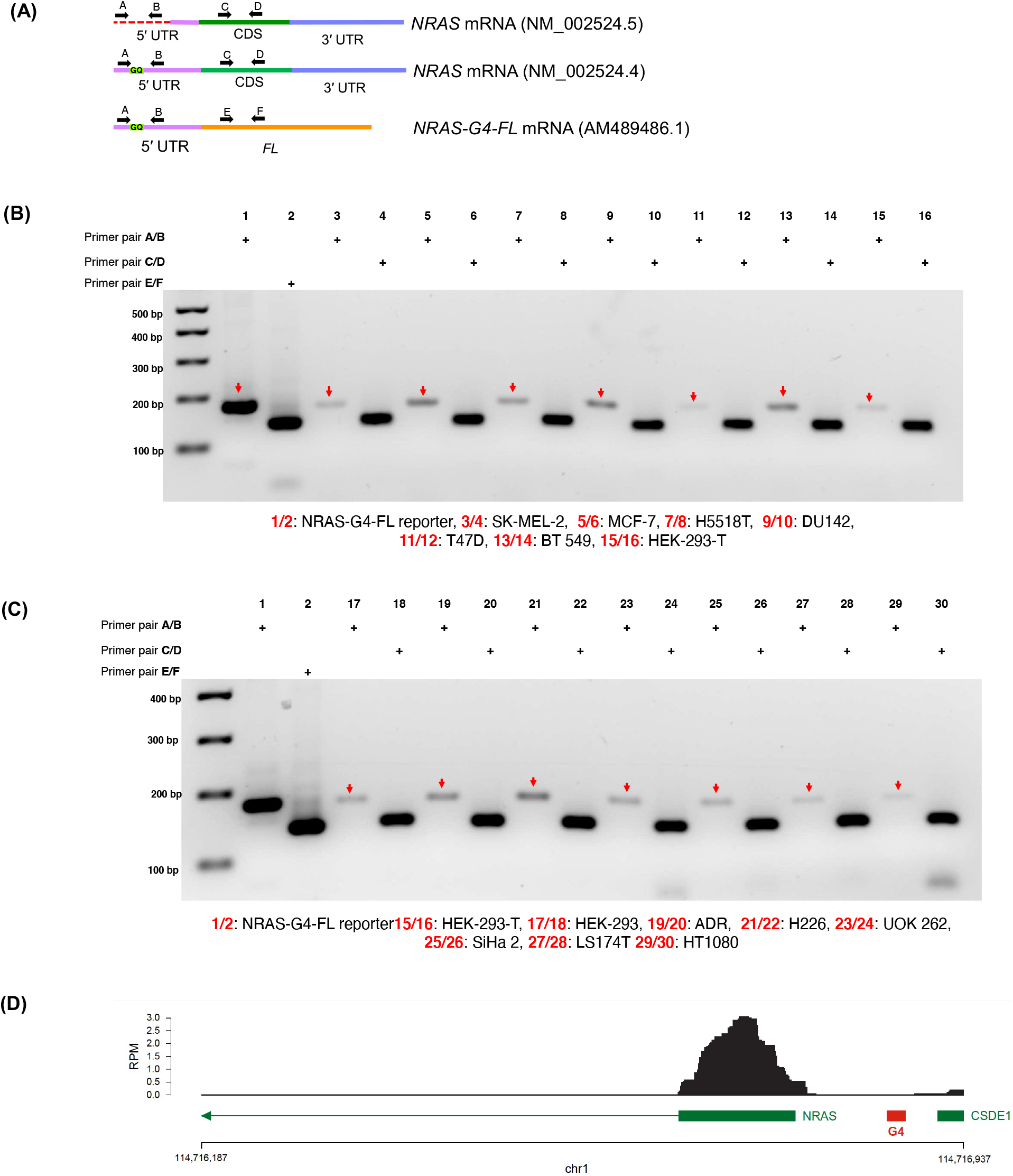
Analysis of the G4 forming region in the 5′ UTR of NRAS. **(A)** Schematic representation of *NRAS* mRNAs (NM_002524.4), (NM_002524.5), and chimeric luciferase construct mRNA (NRAS-G4-FL). The primer binding regions are marked in black arrow (top). The A/B primer pair aligned in the G4 region in the 5′ UTR of *NRAS* mRNA and C/D primer pair aligned in the CDS of NRAS. The E/F primer pair specific for CDS of firefly luciferase mRNA. **(B-C)** The agarose gel electrophoresis of RT-PCR products of NRAS mRNA from 14 different cell lines. Red arrows represent the amplicon of the G4 spanning region in the 5′ UTR of *NRAS* mRNA. **(D)** 5′ RACE analysis of the NRAS TSS in HEK-293 cells. The G4 regions marked with red block and are in the promoter region

To further investigate the existence of G4s in NRAS transcripts in biological contexts, we analyzed genome-wide rG4-seq (G4s in RNA) data reported by the Balasubramanian group^43^. We observed that the G4 forming region in the NRAS transcript has lower mismatch rate in both samples K^+^ and pyridostatin (PDS)-stabilized data sets compared to Li^+^ data set, indicating the lower levels of G4 in the NRAS-5′-UTR (Supplementary Figure S13). For comparison, we analyzed the G4-seq data (G4s in DNA) also reported by Balasubramanian^47^. In this case, a higher mismatch rate was observed in both K+ and PDS-stabilized data sets, indicating that a DNA G4 in the *NRAS* promoter region, upstream of the predominant transcription start site, could be relevant (Supplementary information S14).

## Discussion

Direct targeting of NRAS at the protein level has proven to be highly challenging due to a lack of suitable small molecule binding pockets. Here, we evaluate an alternative approach to control *NRAS* expression via targeting a structured region in the 5′UTR of NRAS mRNA with small molecules. The development of such molecules is an attractive approach to control NRAS expression at the post-transcriptional level. We utilized an SMM screening method to identify small molecule that bind to an rG4 structure reported to be within the *NRAS* 5′UTR. The best compound **18** identified through SAR studies showed reversible binding to the NRAS-G4 structure with submicromolar equilibrium dissociation constants in multiple orthogonal biophysical assays. Further, weaker or no binding was observed in a variety of other RNA and DNA G4 structures. Structural analysis using RNase A probing indicated that the lead compound binds to a site near C12 and C17, which were shown to be on neighboring loops by X-ray crystallography.

The effects of **18** on the translational efficiency of a reporter gene containing the NRAS 5′-UTR were confirmed by *in vitro* translation assays. Compound **18** showed dose-dependent translation inhibition in a wild type luciferase construct (NRAS-G4-FL) but not in a G4-deleted control (NRAS-G4-Del-FL). Using SHAPE-MAP, we demonstrated that the G4 structure folds in the context of this reporter construct (NRAS-G4 -FL). Compound **18** caused an increase in the melting temperature of the G4 in thermal unfolding experiments, confirming that it stabilizes the NRAS-G4 structure upon binding. Taken together, these observations indicate that compound **18** inhibits translation via interactions that stabilize the G4 structure in the NRAS 5′UTR. Thus, this study provides a proof of principle that small molecules that bind to structured elements within the 5′-UTR of mRNAs can block translation, consistent with other studies on the NRAS mRNA.

More in-depth studies aimed at controlling wild type NRAS translation revealed that the compound had only modest effects in cells. RNA seq and qPCR analysis demonstrated no changes in levels of *NRAS* mRNA, and no major changes in a variety of other G4-associated genes. However, the compound had only modest effects on NRAS protein levels in cells, prompting us to investigate the structure of the wild type NRAS transcript. Here, we used a series of qPCR experiments with amplicons designed to cover both the G4 region and coding sequence of the *NRAS* mRNA. Remarkably, we observe that in 14 different cell lines, the predominant NRAS transcript lacks a G4 in the mRNA, and the majority transcription start site is downstream of the G4-forming sequence, though there is some evidence that some transcripts contain the G4 sequence. These qPCR experiments were supported further by 5’ RACE analysis, confirming that the predominant NRAS transcript lacks a G4 in the 5’ UTR in HEK 293 cells. In addition, analysis of multiple public datasets, including CAGE-seq data, rG4-seq, and G4-seq (for DNA G4s) indicate that the G4-forming sequence is mostly not transcribed and may fold into a G4 in the DNA. Finally, it is worth noting that the updated reference sequence for the NRAS mRNA (NM_002524.5) also lacks a G4 in the 5’UTR. Thus, efforts to target NRAS translation with G4 binding small molecules should focus on the identification of cellular contexts in which the G4 is actively transcribed in this gene, or on other regions of the NRAS transcript. Importantly, this work does not rule out the existence of a G4 in NRAS, rather it illuminates an intriguing example where heterogeneity in the expressed transcript impacts the ability to pharmacologically target the RNA. Still, in the broader sense the strategy of targeting rG4 sequences or other highly structured regions in 5’ UTRs with small molecules remains a highly attractive strategy to control mRNA translation with small molecules.

## Supplementary Information

Supporting information is available free of charge at pubs.acs.org. Available data include experimental procedures and compound characterization. RNA sequencing data have been uploaded to the Gene Expression Omnibus (GEO). X-Ray crystallographic datasets have been uploaded to the Protein Databank (PDB).

## Acknowledgments

The authors thank Dr. S. Tarasov and M. Dyba (Biophysics Resource, SBL, NCI at Frederick) for assistance with CD-based experiments. M.T.B is a Lenfant Postdoctoral Fellow of the National Heart, Lung and Blood Institute (NHLBI) This research was supported (in part) by the Intramural Research Programs of the NIH, National Cancer Institute, Center for Cancer Research (grant 1-BC011970) and the NHLBI, NIH. We thank Shankar Balasubramanian for the gift of NRAS luciferase reporter plasmids

