## Supplementary Information for "Investigating the NRAS 5’ UTR as a Target for Small Molecules"

##### Supporting information

### Section 1: Materials and methods

##### SMM screening of a NRAS-GQ:

Small molecule microarray screening was carried out as previously described. Briefly, γ-aminopropyl silane (GAPS) microscope slides (Corning) were functionalized with a short Fmoc-protected amino polyethylene glycol spacer. After deprotection using piperidine, 1,6-diisocyanatohexane was coupled to the surface by urea bond formation to provide functionalized isocyanate-coated microarray slides that can react with primary alcohols and amines to form immobilized chemical screening libraries. A total of 26,227 unique small molecule stock solutions (10 mM in DMSO) from MIPE and NCI Diversity set V screening collections, in addition to dyes and controls, were printed in duplicate onto one slide and exposed to pyridine vapor in a vacuum desiccator overnight to facilitate covalent attachment to the slide surface. After drying, slides were incubated with a 1:20 polyethylene glycol:DMF (v/v) solution to quench unreacted isocyanate surface. The 5'-AlexaFluor647-NRAS-GQ RNA was dissolved in 10 mM Tris buffer (pH 6.9) with 100 mM KCl, diluted to 5μM. Then sample was annealed by heating to 95°C for 3 min, followed by slowly cooling to room temperature for1 hr. The annealed RNA was then further diluted to 1μM in 10 mM Tris buffer (pH 6.9) with 100 mM KCl for screening. Next, printed slides were incubated with the RNA at a concentration of 1μM for 2 hrs at room temperature. Following incubation, slides were gently washed three times for 2 min in 10 mM Tris buffer (pH 6.9) with 100 mM KCl, once in deionized water, and dried by centrifugation for 2 min at 4000 rpm. Fluorescence intensity was measured (650 nm excitation, 670 nm emission) on an Innopsys Innoscan 1100 AL Microarray Scanner with a resolution of 5μm. The scanned image was aligned with the corresponding GenePix Array List (GAL) file to identify individual features. Hits were identified based on signal-to-noise ratio (SNR), defined as (mean foreground − mean background)/standard deviation of background, and Z-score, defined as: Z = (mean SNR635compound – mean SNR635library)/(SD SNR635library) with the following criteria: (i) SNR > 0, (ii) Z score > 3, (iii) coefficient of variance (CV) of replicate spots <100, (iv) SNR of negative control slide <1 and (v) visual comparison of intensity with other nucleic acid structures screened. To further measure selectivity, the Z-score for each selected compound is compared across many different SMM screens (in this case 20 different oligonucleotides, (Supplementary Table S1).

**Water ligand observed gradient spectroscopy (waterLOGSY):**

A reference 1D-$^{1}$H and 1D WaterLOGSY spectrum of 100 μM N-methyl-L-valine and 100 μM compound **1** was collected, followed by a separate sample containing 5 μM NRAS-G4 RNA oligo, 100 μM N-methyl-L-valine, and 100 μM compound **1**. NRAS-G4 RNA (UGUGGGAGGGGCGGGUCUGGG) was buffer exchanged into 10 mM Tris-d11 buffer (pH 6.9, containing 100 mM KCl) using centrifugal filtration (3 kDa MWCO, EMD Millipore) and were annealed by heating to 95 °C for 3 min, followed by slowly cooling to room temperature for 1 hour. A sample of compound **1** and N-methyl-L-valine, each at 100 μM, was prepared in 10 mM Tris-d11 buffer (pH 6.9, containing 100 mM KCl and 5% DMSO-d6), and 1D reference proton and WaterLOGSY spectra with and without oligonucleotide were recorded. These spectra were recorded at 20 °C on a Bruker AVANCE III 500 MHz spectrometer equipped with TCI cryogenically cooled probe. The "zgesgp" excitation sculpting water suppression pulse sequence from Bruker was used for data acquisition with 128 scans. All data were processed and visualized with MestReNova software (Version 8.1.2–11880).

**Thermal melt assays:**

The 10 μM NRAS-G4 RNA was folded in 10 mM Tris (pH=6.9) and 1 mM KCl by heated at 95°C for 3 min and cooled to room temperature over 1 hour. Then 10 μM compound **1** or compound **18** was added into the folded RNA samples and incubated for 15 min in room temperature. For the control sample 5% DMSO was added into the folded RNA sample. Then thermal stability of the NRAS-G4 oligonucleotide with and without compounds was determined by heating from 25 to 97 °C at 1 °C/min in a 0.1 cm quartz cuvette using an Aviv Biomedical Model 420 Circular Dichroism (CD) Spectrometer equipped with a ThermoCube temperature regulator. To calculate the $T_m$ of each sample, ellipticity (folded fraction) was plotted as a function of temperature and fit in GraphPad Prism 7 software using a nonlinear sigmoidal dose-response model with a variable slope. Each condition was performed in triplicate, with $\Delta T_m$ values calculated using $T_m$(+compound) – $T_m$(apo) and then averaged to yield the final value.

**Fluorescence intensity titration:**

AlexaFluor 647-labeled NRAS-G4 RNA or other appropriate oligonucleotide was heated at 95 °C for 3 min, allowed to cool to room temperature for 1 hour, and diluted to 50 nM in 10 mM Tris buffer (pH 6.9, containing 100 mM KCl). Compound was added as a solution both in buffer containing 5% DMSO, and the sample was allowed to equilibrate for 15 min. Fluorescence intensity spectra were recorded at room temperature using a Photon Technology International, Inc. QuantaMaster 600TMSpectrofluorometer equipped with Felix GX 4.2.2 software. Fluorescence intensity was recorded at an excitation wavelength of 645 nm, with the resulting emission spectrum recorded from 650 to 800 nm. Total area under the peak from 655–700 nm was quantified and was then normalized to the values obtained for RNA incubated with a DMSO control. Normalized fluorescence for three independent replicates were averaged and plotted against small molecule concentration. KD values were determined using a single site-binding model.

**Surface plasmon resonance:**

SPR was conducted using a Biacore 3000 (Biacore, Inc) instrument. A CM5 SPR biochip was loaded into the system and primed with running buffer (10 mM Tris, pH 6.9, 100 mM KCl, 0.005% Tween 20, 5% DMSO). For surface making, the flow rate was set as 5 μL/min. Then both Flow Cell (Fc) 1 and 2 were activated by EDC/NHS (0.4 M/0.1 M) aqueous solution for 15 min, followed with an injection of streptavidin (SA) solution (0.2 mg/mL in 10 mM sodium acetate buffer, pH4.5) for 30 min. After the immobilization amount of SA reached 8,000~10,000 RU, the surface was deactivated by flowing 1 M ethanolamine aqueous solution (pH 8.5) for 10 min and regenerated with 10 mM NaOH for 2 min to remove the unbound SA. Then 5 μM folded 5′-biotin NRAS-G4 RNA in 10 mM Tris, 100 mM KCl, pH 6.9 was immobilized on Fc 2 of the SPR chip to a density of 1400 RU. Each of the compound solution was prepared in non-DMSO running buffer by dilution resulting in a final concentration of 5% DMSO. Then, a total of 50 μL compound solution was injected at a flow rate of 30 mL/min in Fc 1-2 flow path for 120 s for association, followed with 200 s running of buffer for dissociation. The final binding curve was obtained by reference subtraction. To determine the binding affinity (KD), a series of diluted compound solutions were injected and KD was calculated by BIAevaluation 4.0 software (GE Healthcare) using Langmuir 1:1 binding model.

**RNAse A Footprinting:**

The 5′-end-AlexaFluor-647-labeled NRAS-G4 RNA was folded in 10 mM Tris (pH=6.9) and 100 mM KCl by heated at 95°C for 3 min and cooled to room temperature over 1 hour. Then increasing concentration of compound **18** (5μM to 100 μM) was added into the folded RNA samples and incubated for 15 min in room temperature. Then the folded RNAs were digested with 0.0001μg of RNase A (Ambion) for 3 min at room temperature. The reactions were terminated by heating at 95 °C for 5 min with an equal volume of stop buffer (7 M urea, 10 mM Tris-HCl (pH 7.5), and 0.1 mM EDTA). Samples were electrophoresed on a 17% denaturing PAGE. The gel images were obtained by scanning the screen on a Typhoon Imager (Amersham) and bands were quantified by using Image J software.

***In vitro* Transcription:**

The plasmids pSKC11 and pSKC12, which encode the transcripts NRAS-G4-FL and NRAS-G4-Del-FL, respectively were received from Balasubramanian group.[1] Then the plasmids were linearized at the 3′ end using EcoRI, and linearized plasmids were used as a template for *invitro* transcription. The 5′-capped transcripts were synthesized *in vitro* using mMessage mMachine T7(ThermoFisher Scientific), followed by incubated with DNase for 15minutes at 37 °C to remove the template DNA. Then all the transcripts were purified using Monarch RNA Cleanup Kit (NEB). The concentration was determined using a Nanodrop. Integrity and size of each transcript was confirmed using a 1% agarose gel.

***In vitro* translation and luciferase assay:**

*In vitro* translation of NRAS-G4-FL and NRAS-G4-Del-FL transcripts in the presence of compound **18** were carried out in a cell-free translation system of rabbit reticulocyte lysates (Promega) following manufacturer’s protocol. Briefly, 500 ng of NRAS-G4-FL and NRAS-G4-Del-FL transcripts were folded by heated at 95°C for 5 min and slowly cooled down to room temperature for 1 hour. Then compound **18** (1 μM, 10 μM and 20 μM) or 5% DMSO were added into the folded RNA and incubated for 15 minutes at room temperature. The *in vitro* translation was carried out by adding the reticulocyte lysate into the samples and incubated at 30°C for 90 min. The firefly luciferase activity was measured using luciferase assay reagent (Promega) according to manufacturer’s protocol on a Synergy Mx microplate reader (BioTek).

**In vitro RNA structure probing:**

*In vitro* RNA folding and probing was carried out as previously described[2]. Briefly, 2 μg of NRAS-G4-FL RNA in 89ul nuclease free $H_2O$ was heated at 95°C for 2 min, then immediately transferred to ice for 1 min. Then10 μl of ice-cold RNA Folding Buffer 10X (500 mM HEPES pH 7.5; 1 M KCl) were then added, the solution mixed, and incubated at 37°C for 15 min. Then, 1 μl of 1 M $MgCl_2$, pre-warmed at 37°C, was added, the solution mixed, and further incubated for 15 min. Probing was performed by adding 11 μl of NAI (1M stock), incubating at 37°C with moderate shaking for 10 min. For the control reaction, 11μl of neat DMSO was added. Reaction was quenched by adding 111μl of 1M DTT. RNA was recovered by purification on a Monarch RNA 10μg column (New England Biolabs).

**Library preparation:**

Both DMSO- and NAI-treated samples, were fragmented in a buffer containing 4mM final $MgCl_2$, by incubating at 94°C for 8 min. The fragmented RNA was then subjected to random-primed reverse transcription, as per standard SHAPE-MaP conditions. Briefly, fragmented RNA was mixed with 1μl of 10μM random decamers, and 1μl 10mM dNTPs, then incubated at 70°C for 5 min, and immediately transferred to ice for 1 min. Then, 4 μl of 5X RT Buffer (250mM Tris pH 8.0; 375mM KCl), 2 μl of DTT (0.1M), 1 μl of SuperaseIN RNase inhibitor (Ambion), 1 μl of SuperScript II RT (ThermoFisher Scientific), and 1 μl 120mM $MnCl_2$ were added, the reactions mixed, and incubated at 25°C for 10 min, followed by 2 hours at 42°C. After reaction cleanup, the buffer was replaced with the standard SuperScript II RT First Strand Buffer with $MgCl_2$, and the RNA-cDNA hybrids were used as input for the NEBNext® Ultra™ II Non-Directional RNA Second Strand Synthesis Module (New England Biolabs). After having converted the RNA into dsDNA, the remainder of the library preparation was then carried out using the NEBNext® Ultra™ II DNA Library Prep Kit for Illumina® (New England Biolabs), as per manufacturer instructions.

**Data analysis:**

All the relevant analysis steps have been conducted using the RNA Framework (https://github.com/dincarnato/RNAFramework)[3]. SHAPE raw reactivities were calculated via the rf-norm module, by using the previously published method[4], followed by box-plot normalization.

**Crystallization and Structural Determination:**

The NRAS-G4 variant consisting of a G8U mutation (NRAS-G8U) was chemically synthesized by Dharmacon and purified using denaturing polyacrylamide gel electrophoresis. The NRAS-G8U construct used for crystallization comprised of 22 nucleotides (5´-UGUGGGAUGGGCGGGUCUGGGA-3´). Prior to crystallization, the NRAS-G8U was heated to 95°C for 1.5 minute and allowed to cool overnight in 50 mM potassium chloride and 25 mM sodium cacodylate at pH 6.8. Crystals of NRAS-G8U were produced using the vapor diffusion method at 21°C by combining 1 µL each of the folded RNA and reservoir solution (20 - 25 % PEG 3350, 80 mM sodium chloride and 100 mM BIS-TRIS pH 6.5). After two months, small tetragonal bipyramidal crystals (50 x 50 x 150 µm) were supplemented with 30 % PEG 3350 and plunged into liquid nitrogen. X-ray diffraction data was collected at a wavelength of 1.104 Å at beamline 17-ID-C located at the Advance Photon Source (Argonne National Laboratory), and reduced using the XDS suite[5], AIMLESS[6] and POINTLESS[7]. The NRAS-G4 structure was solved by molecular replacement[8] using a NMR solution structure of a parallel DNA G4[9] (PDB: 5NYS) as a search model (Top LLG = 74.4, Top TFZ = 8.7). The final model was produced from iterative cycles of rebuilding in COOT[10] with interspersed refinements using Phenix.refine[11]. The following refinement strategy was implemented: torsion-angle simulated annealing, XYZ refinement, and individual isotropic B-factor. Crystallographic and refinement statistics for NRAS-G8U can be found in Table S5. The NRAS-G8U structure was deposited in the PDB possessing the accession code 7SXP.

**Cell culture**

MCF-7 cells and SK-MEL-2 cells were grown in Dulbecco's modified Eagle's medium (DMEM) with 1% glutamine, 10% fetal bovine serum and 1% antibiotics (streptomycin and penicillin) at 37°C in 5% CO2 in a humidified incubator according to ATCC's recommendations. Cells were grown in 96 well plates (for MTS assay) or 6-well plates for (RT-PCR, qPCR and western blotting).

**Cell Viability Assay:**

MCF-7 and SKE-MEL-2 cells were plated in 96-well culture plates at a density of $1 \times 10^4$ cells/mL for MCF-7 cells and $2.5 \times 10^4$ cells/mL for SK-MEL-2 cells. After 24-hour incubation, varying concentration of the compound **18** or DMSO (control) were added to the cells and were further incubated for 72 hours. Following the aforementioned incubation time, 3-(4,5-dimethyl thiazol-2-yl)-2,5-diphenyltetrazolium bromide (MTT) was directly added into the wells and incubated again at 37 °C with 5% CO2 for 4 hours. The ability of cells to form formazan crystals by active mitochondrial respiration was determined using a Microplate reader at 540 nm after dissolving the crystals in an SDS-HCl solution. A blank measurement was taken from the absorbance of the wells with media only and subtracted accordingly. The cell viability percentages were calculated by normalizing to the untreated control cells. The $IC_{50}$ (inhibitory concentration to produce 50% cell death) values were determined by fitting the data in dose−response curves in prism. Data were presented as the mean ± SEM.

**qPCR:**

MCF-7 and SKE-MEL-2 cells were seeded in 6-well culture plates at a density of 1 × $10^6$ cells/mL for MCF-7 cells and 2.5 × $10^6$ cells/mL for SK-MEL-2 cells. After 24-hour incubation, different doses of the compound **18** (1, 5, 10 and 25 µM) or DMSO (control) were added to the cells and were further incubated for 48 hours. Then total RNAs were isolated using E.Z.N.A.® MicroElute Total RNA Kit (Omega), and DNA was removed by on-membrane DNase I (Omega) digestion according to the manufacture's protocol. Then 1 µg of RNA was used for cDNA synthesis. The cDNA was synthesized using qScript cDNA SuperMix (Quanta Biosciences) and oligo d(T) primer according to the manufacture’s protocol. The 20 µl reactions were incubated in a thermocycler (MiniAmp plus Applied Biosystem) for 5 min at 22 °C, 30 min at 42 °C, 5 min at 85 °C, then held at 4 °C. Then 60 ng of cDNAs were subjected to qPCR using a Perfecta SYBR Green Super Mix (Quanta Biosciences) on an Eppendorf Mastercycler RealPlex2 in the presence of appropriate set of primers. The reactions were incubated at 95 °C for 10 min, followed by 40 cycles of 95 °C for 30 sec, 60 °C for 30 sec and 72 °C for 20 sec. Threshold cycles (CT) is the first cycle that showed a detectable increase in fluorescence due to the formation of PCR products and was used to determine the template amount in each sample. The relative fold change in expression was measured using the Livak method and were normalized to the relative level of mRNAs of the control experiment[12]. For example, to calculate the ΔΔ(CT) between each gene of interest and the average of the control samples: Δ(CT) = CT (NRAS) − CT(GAPDH); ΔΔ(CT) = ΔCT (treatment) − ΔCT (control); fold change = $2^{-\Delta\Delta\ (CT)}$. The primers used are shown in the supplementary information Table S6.

**Western blotting:**

MCF-7 and SKE-MEL-2 cells were seeded in 6-well culture plates at a density of $1 \times 10^6$ cells/mL for MCF-7 cells and $2.5 \times 10^6$ cells/mL for SK-MEL-2 cells. After 24-hour incubation, different doses of the compound **18** (1, 5, 10 and 25 μM) or DMSO (control) were added to the cells and were further incubated for 48 hours. Proteins were extracted from the cells with RIPA buffer (RIPA, sodium orthovanadate, PMSF, protease inhibitor, and phosphatase inhibitors A and B), vortexed to homogenize, and sonicated with intervals of 1s on, 30 seconds off, for 1 minute. The amount of protein was quantified by a standard Bradford protocol. Then, 35 μg of protein was loaded into each well of 4–12% Bis-Tris Gels (Novex), electrophoresed at 180 V for 60 min. The Ponceau staining (Thermo Scientific) was performed to confirm equal loading and transfer. Then blots were blocked with 1x Blocking buffer (thermo Scientific) for 1 hour, washed three times in

1XTBST for 10 min each. The NRAS and GAPDH proteins were detected by incubating with NRAS (abcam ab-188369) antibody and GAPDH (ThermoFisher MA5-15738) antibody at 4 °C overnight. Horseradish peroxidase-conjugated rabbit anti-mouse IgG (abcam ab-6728) was used as the secondary antibody at 1:1000 dilutions for GAPDH and NRAS. Proteins were visualized by Western Blotting Luminol Reagent (Cell Signaling Technology 7003) in Image Quant LAS 4000 (GE healthcare).

**RNA Seq:**

MCF-7 and SKE-MEL-2 cells were seeded in 6-well culture plates at a density of 1 × $10^6$ cells/mL for MCF-7 cells and 2.5 × $10^6$ cells/mL for SK-MEL-2 cells. After 24-hour incubation, 1 μM compound **18** or 1% DMSO (control) were added to the cells and were further incubated for 48 hours. Then total RNAs were isolated using E.Z.N.A.® MicroElute Total RNA Kit (Omega), and DNA was removed by on-membrane DNase I (Omega) digestion according to the manufacture's protocol. Then, isolated RNA integrity and quality were determined by Bioanalyzer (Agilent 2100) by using the Agilent RNA 6000 Nano kit. Then, samples were submitted to Sequencing facility (CCR-SF). Then, 500 ng of total RNA was used as the input for mRNA capture with oligo-dT coated magnetic beads. The library was prepared using Illumina TruSeq Stranded mRNA Library kit according to manufactured protocol and final purified products were then quantitated by qPCR before cluster generation and sequencing. Then, samples were sequenced 2x 76 cycles run on NextSeq500 sequencer. The samples have 21 to 56 million pass filter reads with more than 94% of bases above the quality score of Q30.

**RNA-Seq Data Analysis:**

Upon generation of FASTQ files by CCR-SF, the sequencing analysis was performed using the Center for Cancer Research Collaborative Bioinformatics Resource (CCBR) RNA-Seq pipeline (https://github.com/CCBR/Pipeliner). The CCBR pipeline was then deployed on the National Institutes of Health (NIH) HPC Biowulf cluster (https://hpc.nih.gov). In brief, the CCBR pipeline trims the sequencing adapters using the Cutadapt tool before alignment with the reference genome (Human - hg38) using STAR's two-pass alignment system. The mapping statistics were calculated using Picard software. The average mapping rate of all samples was 95% and unique alignment was above 86%. There were 3.49 to 7.70% unmapped reads, and the samples have 0.20% ribosomal bases. Percent coding bases were between 56-61%, percent UTR bases were 32-36%, and mRNA bases were between 90-94% for all the samples. Library complexity was measured in terms of unique fragments in the mapped reads using Picard's MarkDuplicate utility. The samples have 83-87% non-duplicate reads. Then, gene expression quantification analysis was performed using the RSEM tool yielding raw and normalized counts. Upon generation of gene counts by RSEM, a CPM (counts per million) filter (CPM < 0.5) was used to remove lowly expressed genes. Then, DESeq2 package was used to perform comparative analysis and derive fold change in expression and false discovery rate (FDR) adjusted p-values needed for generation of volcano plots. In addition to the output from the CCBR pipeline, we applied an additional Transcripts Per Kilobase Million (TPM) filter to only keep genes that have TPM > 3 in more than two samples when comparing an experimental set to DMSO controls. Only genes passing this TPM filter were shown in the volcano plots and used for gene ontology (GO) and pathway analysis.

**RNA-Seq Pathway and GO Enrichment Analysis:**

Enrichment analysis on pathways and GO of significantly differentially expressed genes was performed using Enrichr (maayanlab.cloud/Enrichr/). Significantly upregulated genes were identified as genes with fold change > 1.5 comparing to DMSO with FDR < 0.05. Significantly downregulated genes were chosen using fold change < -1.5 comparing to DMSO with FDR < 0.05. Significantly differentially expressed genes were then fed to Enrichr and "KEGG 2021 Human" was chosen under the "Pathways" tab for the pathway enrichment analysis and "GO Biological Process 2021" was chosen under the "Ontologies" tab for GO analysis. Genes with highest ranked Enrichr "Combined score" were explored for further analysis.

**GEO accession codes: GSE191144**

**RT-PCR:**

All the cell lines (MCF-7, SKE-MEL-2, SiHa, H5518T, DU142, T47D, Bt 549, HEK-293T, HEK-293, ADR, H226, UOK 262, LS174T and HT1080) were grown in T75 flask with proper medium according to the ATCC protocol. Once the cells reached 80% confluence, total RNAs were isolated using TRIzol reagent (Thermofisher scientific), and DNA was removed by TURBO DNase (Thermofisher scientific) digestion according to the manufacture's protocol. Then first strand cDNA was synthesized by Superscript first-strand synthesis kit (Invitrogen) with oligodT primers according to the manufactured protocol. Briefly, 1ug of RNA and oligodT were mixed together and heated at 65°C for

5min to denature the RNA. Then samples were placed in ice for 10 min. In a separate tube, 2X reaction mix was prepared by adding the 2μl of 10X RT buffer (containing the LiCl), 4μl of 25mM $MgCl_2$, 2μl of 0.1M DTT and 1μl of RNaseOUT. The 9μl of reaction mix was added into the RNA/primer mix samples and incubated at 42°C for 2min. Then 1μl of SuperScript II RT was added into the samples and incubated at 42 °C for 50 min, followed by incubated at 70 °C for 15 min to heat inactivate the enzyme. Then 2μl of cDNA was used for PCR amplification with Platinum SuperFi II DNA polymerase (Invitrogen). The reaction was incubated at 98 °C for 30 sec, followed by 25 cycles of 98 °C for 10 sec, 63 °C for 35 sec and 72 °C for 30 sec. Then samples were loaded into the 2% agarose gel and run for 1hour. The specific set of primers that were used to amplify the 5′ UTR and CDS of the NRAS mRNA and CDS of NRAS-G4-FL reporter construct mRNA have shown in the supplementary information Table S9.

**5′-RACE**

To map TSSs, 2 μg of polyA RNA from HEK293 cells were fragmented to an average size of 200 nt by incubating at 94°C for 5 min in the presence of 4mM MgCl2. Endogenous 5′P sites and 2′-3′ cyclic P sites were resolved by treatment with 1U rSAP (New England Biolabs) in CutSmart buffer at 37°C for 30 min. Capped RNA fragments were then decapped using 5U Cap-Clip Acid Pyrophosphatase (Cellscript) at 37°C for 1 hour. Treatment with Cap-Clip leaves a 5′P that can be exploited for direct adaptor ligation. Hence, only capped fragments will be ligated to both a 5′ and a 3′ adapter, and as such, amplified. RNA fragments were then used as input for the NEBNext® Multiplex Small RNA Library Prep Set for Illumina (New England Biolabs).

#### Section 2: Figures and Tables

**Supplementary Table S1:** Selected sequences of RNA and DNA oligonucleotides screened by SMM

| **Name** | **Structure** | **Sequence (5′ -3′)** |
|---|---|---|
| NRAS | RNA-G4 | AlexaFluor647*−UGUGGGAGGGGCGGGUCUGGG |
| TERRA | RNA-G4 | AlexaFluor647*−GGGUUAGGGU |
| ZiKa 3′UTR | RNA-G4 | AlexaFluor647*−GCGGCGGCCGGUGUGGGGAA |
| EWSR1 | RNA-G4 | AlexaFluor647*−GGGGCAGGGGAAGAGGGGG |
| AKTIP | RNA-G4 | AlexaFluor647*−GGGGUGGGGCGGGGCGGG |
| MYB | DNA-G4 | AlexaFluor647*−GGAGGAGGAGGTCACGGAGGAGGAGGAGAAGGAGGAG GAGGA |
| MYC | DNA-G4 | AlexaFluor647*−AGGGTGGGGAGGGTGGGG |
| KRAS | DNA-G4 | AlexaFluor647*−AGGGCGGTGTGGGAAGAGGG AAGAGGGGGAGGCA |
| RB1 | DNA-G4 | AlexaFluor647*−CGGGGGGTTTTGGGCGG |
| BCL2 | DNA-G4 | AlexaFluor647*−AGGGGCGGGCGCGGGAGGAA GGGGGCGGGA |
| cKIT | DNA-G4 | AlexaFluor647*− AGGGAGGGCGCTGGGAGGAGGG |
| FGFR | RNA 3-way junction | AlexaFluor647*−GCUCUUGCUUCGCUUGUUUUCUAGGCCGCCGGUGUUA ACACCACGGACAAAGAGC |
| MALAT1 | RNA triple helix | AlexaFluor647*− AAAGGUUUUUCUUUUCCUGAGAAAUUUCUCAGGUU UUGCUUUUUAAAAAAAAAGCAAAA |
| PreQ$_1$-RS(*BS*) | Riboswitch | AlexaFluor647*− AGAGGUUCUAGCUACACCCUCUAUAAAAAACUAA |
| SAMII-RS | Riboswitch | Cy5*-UCGCGCUGAUUUAACCGUAUUGCAAGCGCGUGAUAAAUGUAGCU AAAAAGGG |
| TPP-RS | Riboswitch | Cy5*−CAGUACUCGGGGUGCCCUUCUGCGUGAAGGCUGAGAAAUACCCG UAUCACCUGAUCUGGAUAAUGCCAGCGUAGGGAAGUGCUG |
| Pre-miR-21 | RNA hairpin | AlexaFluor647*−UAGCUUAUCAGACUGAUGUUGACUGUUGAAUCUCAUGG CAACACCAGUCGAUGGG CUGUC |
| Pre-miR-17 | RNA hairpin | AlexaFluor647*− CAAAGUGCUUACAGUGCAGGUAGUGAUAUGUGCAU CUACUGCAGUGAAGGCACUUGUAG |
| Pre-miR-31 | RNA hairpin | AlexaFluor647*− AGGCAAGAUGCUGGCAUAGCUGUUGAACUGGGAAC CUGCUAUGCCAACAUAUUGCCAUC |
| HIV RRE IIB | RNA hairpin | AlexaFluor647*−UGGGCGCAGUGUCAUUGACG CUGACGGUACA |

*5′-Modification accomplished with AlexaFluor647-NHS ester or Cy5-NHS ester

**Supplementary Table S2:** Commercial Supplier IDs for 30 candidate compounds from the SMM.

| **Supplier Product ID** | | |
|---|---|---|
| C660-1103 (**1**) | 5941-1312 | 14924624 |
| 21964951 (**S1**) | D718-0730 | 35026530 |
| 15888495 (**S2**) | C769-1975 | |
| 20991445 (**S3**) | 27260860 | |
| 15973392 (**S4**) | 18903993 | |
| 13312934 (**S5**) | 8006-4483 | |
| 58902615 (**S6**) | 3702-1289 | |
| 15043612 (**S7**) | 20062634 | |
| 5607-0251 (**S8**) | 17301753 | |
| 32415151 (**S9**) | 17766103 | |
| 42254132 (**S10**) | 3261-0434 | |
| C301-8687 (**S11**) | 2435-0117 | |
| 6359-0041 (**S12**) | 4229-0068 | |
| 31891223 (**S13**) | C769-1518 | |

* IDs with hyphenated numbers are available from ChemDiv (www.chemdiv.com) and non-hyphenated numbers are from ChemBridge (www.chembridge.com).

**Supplementary Table S3:** Selectivity of 14 purchased hit compounds screened by SMM as determined by Z-score and the $K_D$ values determined by SPR assay.

| Supplier Product ID | Compound | Z-Score | SPR KD ( μM) |
|---|---|---|---|
| C660-1103 | **1** | 7.40 | 0.45 ± 0.11 |
| 21964951 | **S1** | 10.81 | 10.9 ± 1.2 |
| 15888495 | **S2** | 3.12 | 25.4 ± 9.3 |
| 20991445 | **S3** | 4.30 | 14.4 ± 10.3 |
| 15973392 | **S4** | 3.10 | 27.9 ± 4.8 |
| 13312934 | **S5** | 4.52 | 39.0 ± 12.6 |
| 58902615 | **S6** | 4.60 | 93 ± 15.3 |
| 15043612 | **S7** | 3.35 | 96 ± 9.7 |
| 5607-0251 | **S8** | 3.20 | > 100 |
| 32415151 | **S9** | 3.47 | 48.1 ± 3.5 |
| 42254132 | **S10** | 3.6943 | N.D |
| C301-8687 | **S11** | 3.24 | > 100 |
| 6359-0041 | **S12** | 12.27 | > 100 |
| 31891223 | **S13** | 17.94 | N.D |

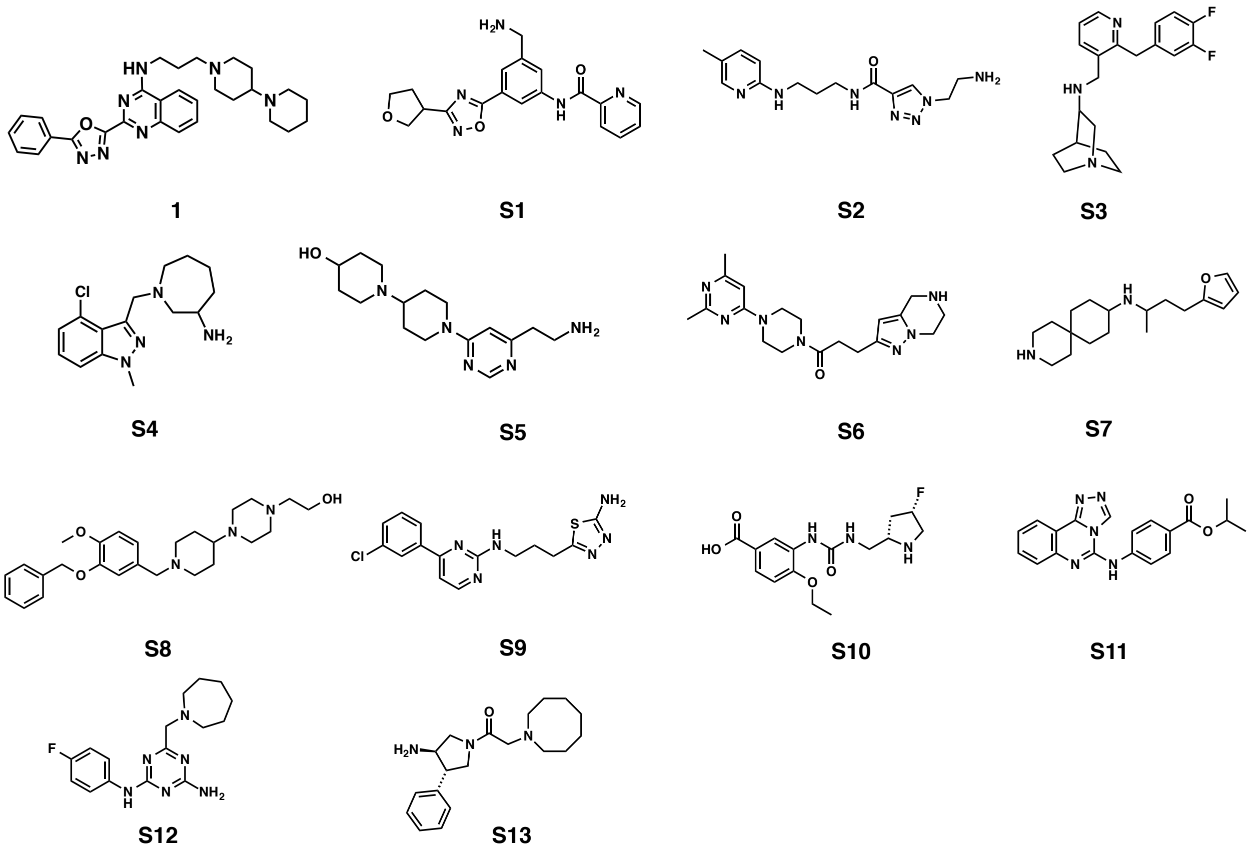


**Figure S1:** Structures of 14 hit compounds for NRAS-G4 identified by SMM screening that were purchased for further analysis.

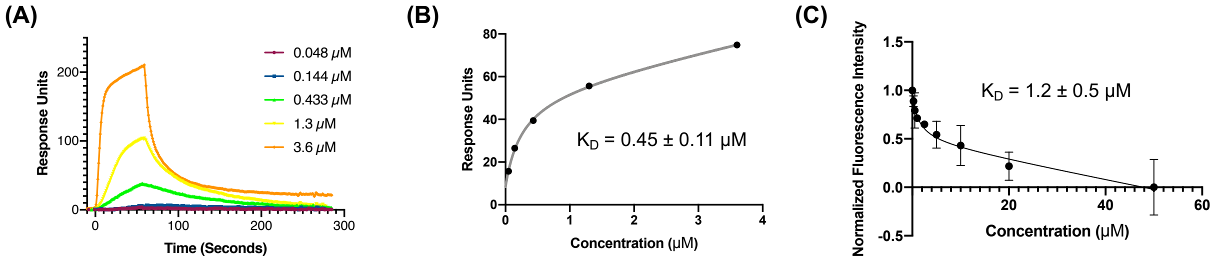


**Figure S2: (A)** Sensorgrams corresponding to 0.048 to 3.8 μM injections of compound **1** over a 5′ biotin labeled NRAS-G4 RNA in SPR. **(B)** The binding curve generated by fitting the response units against the concentration of compound compound **1**. **(C)** Fluorescence intensity assay of 5′-AlexaFluor 647-labeled NRAS-G4 RNA in the presence of increasing concentration of compound **1**. Error bars indicate the standard deviation determined from three independent measurements.

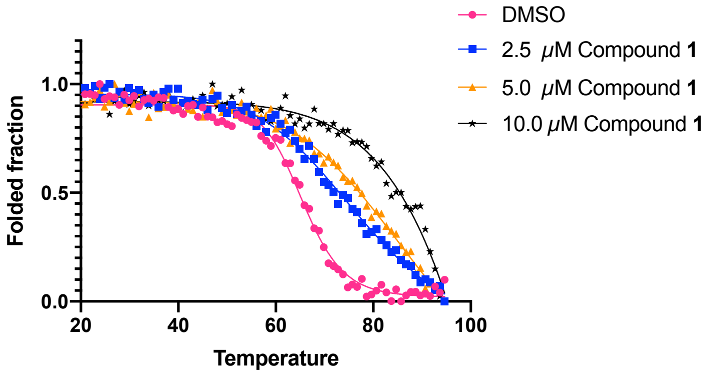


| [ **1** ] (μM) | $T_m$ |
|---|---|
| 0 | 64. 8 ± 1.9 |
| 2.5 | 71.7 ± 1.3 |
| 5.0 | 77.6 ± 1.5 |
| 10.0 | 84.7 ± 2.3 |

**Figure S3:** Circular Dichroism based thermal melting of NRAS-G4 in the absence and presence of increasing concentration of compound **1** (left). The calculated $T_m$ values were indicated in the table (right).

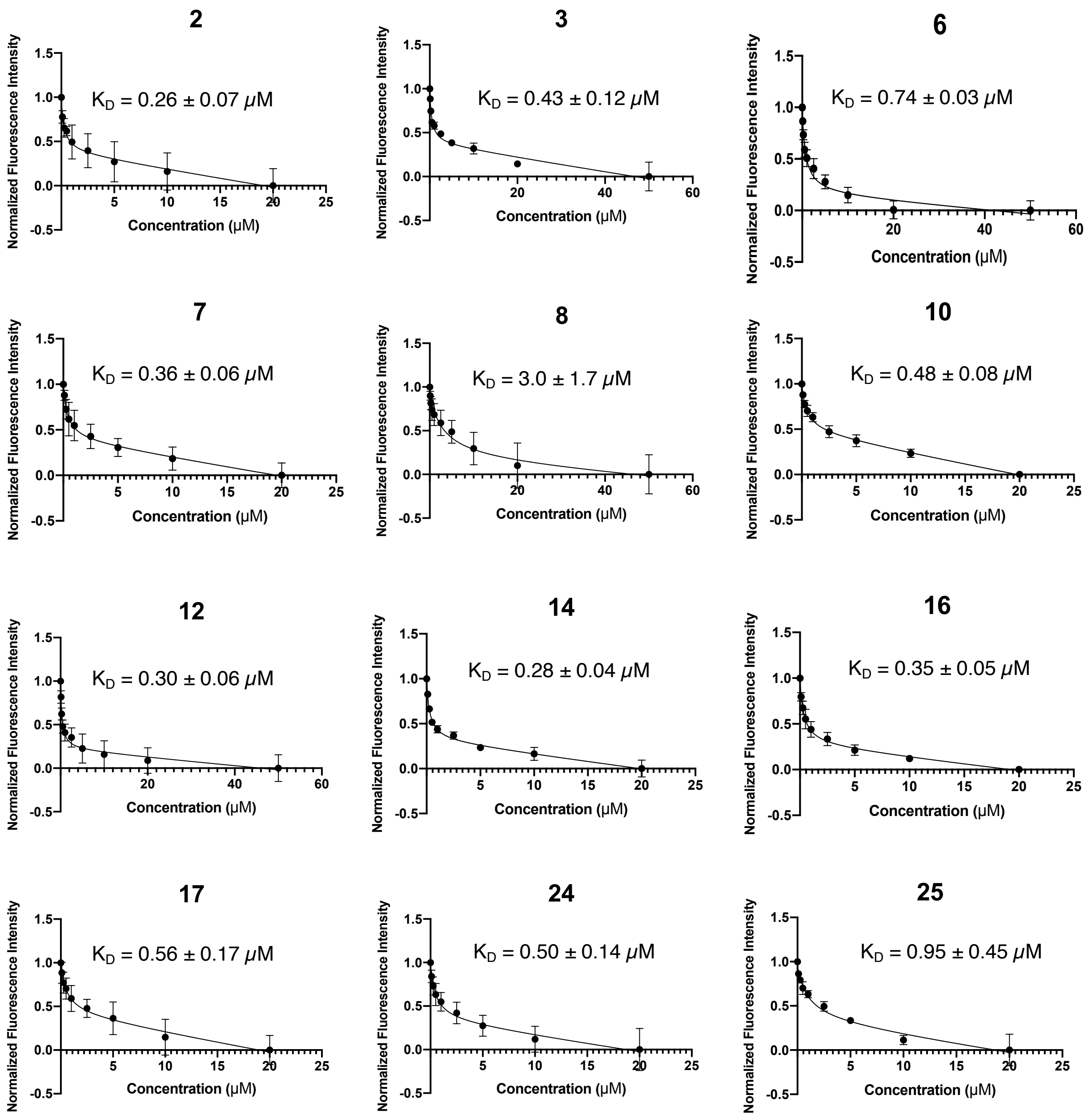


**Figure S4:** Structure-activity relationship (SAR) study of compound **1**. Binding assays were performed by Fluorescence intensity assay. Error bars indicate the standard deviation determined from three independent measurements.

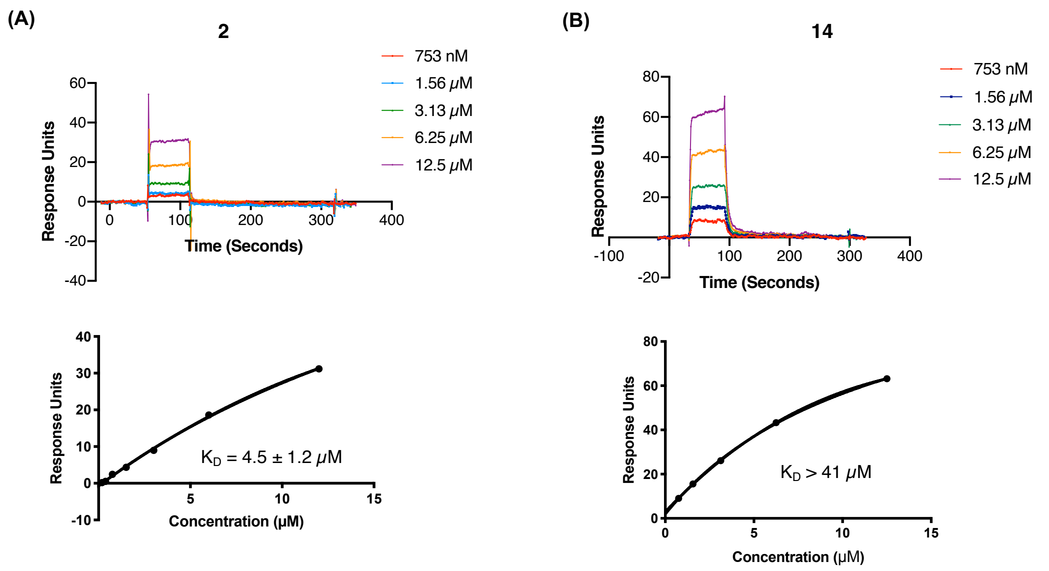


**Figure S5**: **(A)** Sensorgrams corresponding to 753 nM to 12.5 μM injections of compound **2** over a 5′ biotin labeled NRAS-G4 RNA in SPR (top) and the binding curve generated by fitting the response units against the concentration of compound compound **2** (bottom). **(B)** Sensorgrams corresponding to 753 nM to 12.5 μM injections of compound **12** over a 5′ biotin labeled NRAS-G4 RNA in SPR (top) and the binding curve generated by fitting the response units against the concentration of compound compound **12** (bottom).

**Table S4:** G-quadruplex sequences used for binding selectivity profiling.

| Name | Sequence (5′ to 3′) |
|---|---|
| BCL2 DNA G4 | AGG GGC GGG CGC GGG AGG AAG GGG GCG GGA |
| KRAS DNA G4 | AGG GCG GTG TGG GAA GAG GGA AGA GGG GGA GGC AG |
| mTOR DNA G4 | GGG GAA GGC GGG CGG TGG GGC AGG GGG |
| Telomeric DNA G4 | TTA GGG TTA GGG TTA GGG TTA GGG TTA |
| VEGF DNA G4 | CGG GGC GGG CCG GGG GCG GGG T |
| MYCN DNA G4 | AGG GGG TGG GAG GGG GCA TGC AGA TGC AGG GGG |
| AKTIP RNA G4 | GGG GUG GGG CGG GGC GGG |
| TERRA RNA G4 | GGG UUA GGG U |
| EWSR1 RNA G4 | GUG GGC GGG GAG GAG GAC GCG GCG GUG GAA U |

*All the sequence has 5′ end fluorophore label (Cy5 or AlexaFluor647)

**Table S5**: NRAS-G4 crystallographic and refinement statistics (PDB code: 7SXP).

| | NRAS-G4 |
|---|---|
| **Data Collection** | |
| Space Group | P $3_1$21 |
| Cell Dimensions | |
| $a, b, c$ (Å) | 36.4, 36.4, 70.1 |
| $\alpha, \beta, \gamma$ (°) | 90, 90, 120 |
| Resolution (Å) | 70.1 – 2.32 (2.42 – 2.32) |
| $R_{merge}$ | 0.218 (>1.0) |
| $\langle I \rangle / \langle \sigma(I) \rangle$ | 5.3 (1.2) |
| Completeness (%) | 99.6 (99.2) |
| Redundancy | 17.4 |
| **Refinement** | |
| Resolution (Å) | 31.5 – 2.9 (3.0 – 2.9) |
| No. Reflections | 1331 (135) |
| $R_{work}$ / $R_{free}$ | 24.6 / 26.7 |
| No. Atoms | 426 |
| RNA | 413 |
| Ligands | 13 |
| $B$-factors (Å$^2$) | 90.7 |
| RNA | |
| Ligands | 112.2 |
| R.M.S. Deviations | |
| Bond lengths (Å) | 0.007 |
| Bond angles (°) | 1.06 |

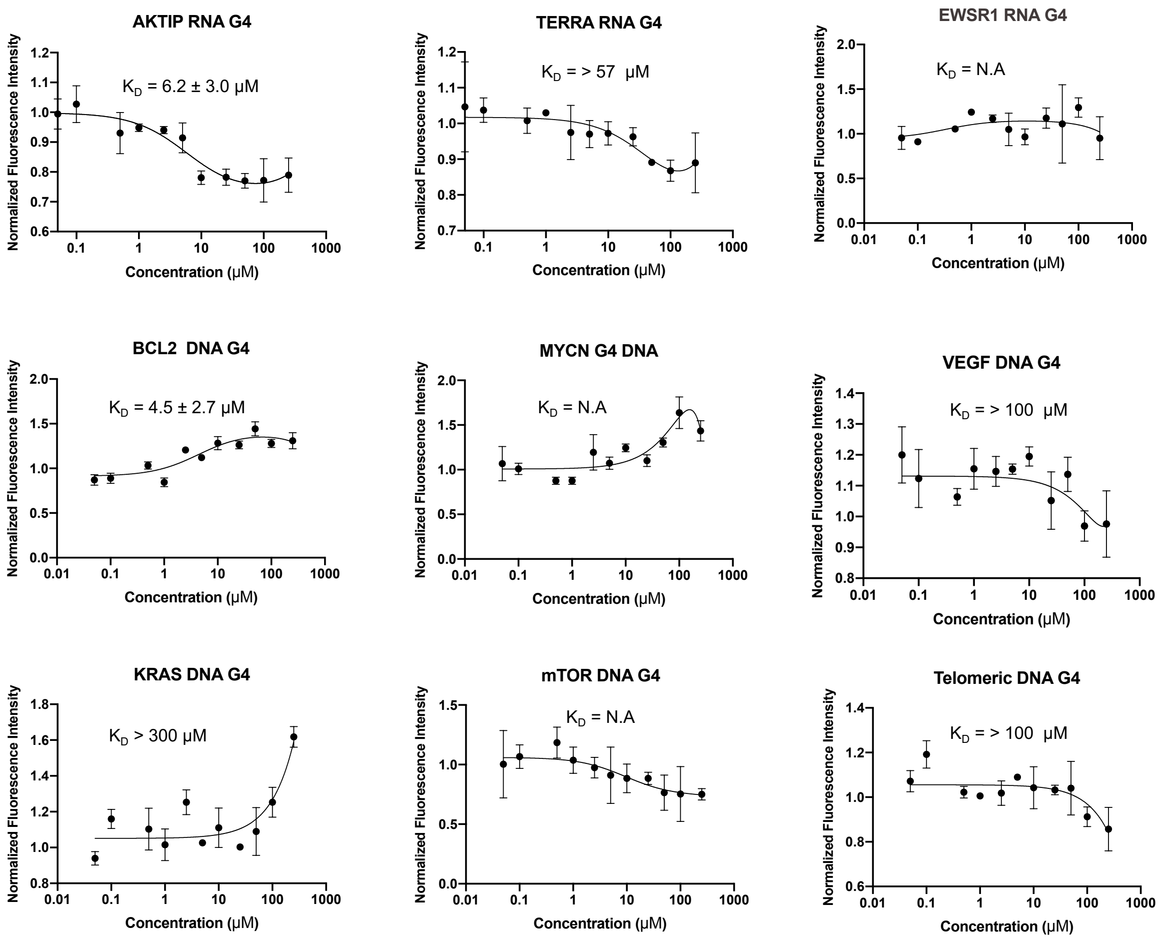


**Figure S6:** Binding affinity determination of compound **18** with nine different G4 targets. All the DNA/RNA G4 oligos were labeled with fluorophore at 5′ end. Titration curves were obtained by fluorescence intensity assay. Error bars indicate the standard deviation determined from three independent measurements.

**Table S6:** The sequence information of the primers used in the qPCR.

| **Name** | **Sequence (5′ to 3′)** |
|---|---|
| GAPDH FP | GCA CCG TCA AGG CTG AGA AC |
| GAPDH RP | TGG TGA AGA CGC CAG TGG A |
| NRAS FP | CCT ATA CAA TGT ATG TAA TTT GTT TCC |
| NRAS RP | CAA TGC ACC AAA GTT TTA CAA TAT TTG AAC |

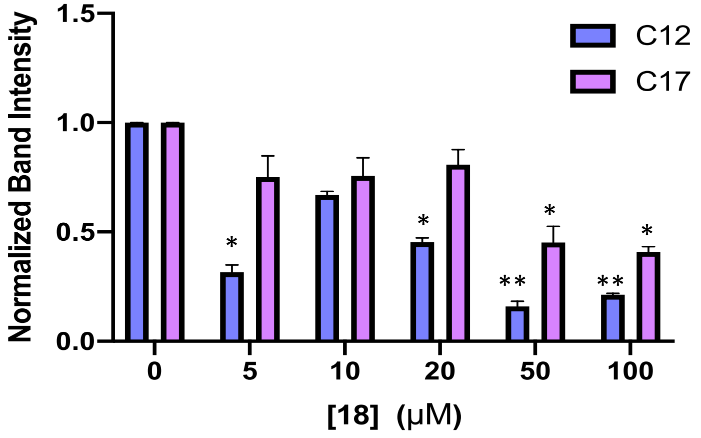


**Figure S7:** The bar graph represents the quantification of band intensity of C12 and C17 with respect to concentration of **18**, where band intensity of C12 and C17 from samples treated with **18** was normalized to the average band intensity of DMSO control treated samples. (n=3, **P < 0.01 and *P < 0.05).

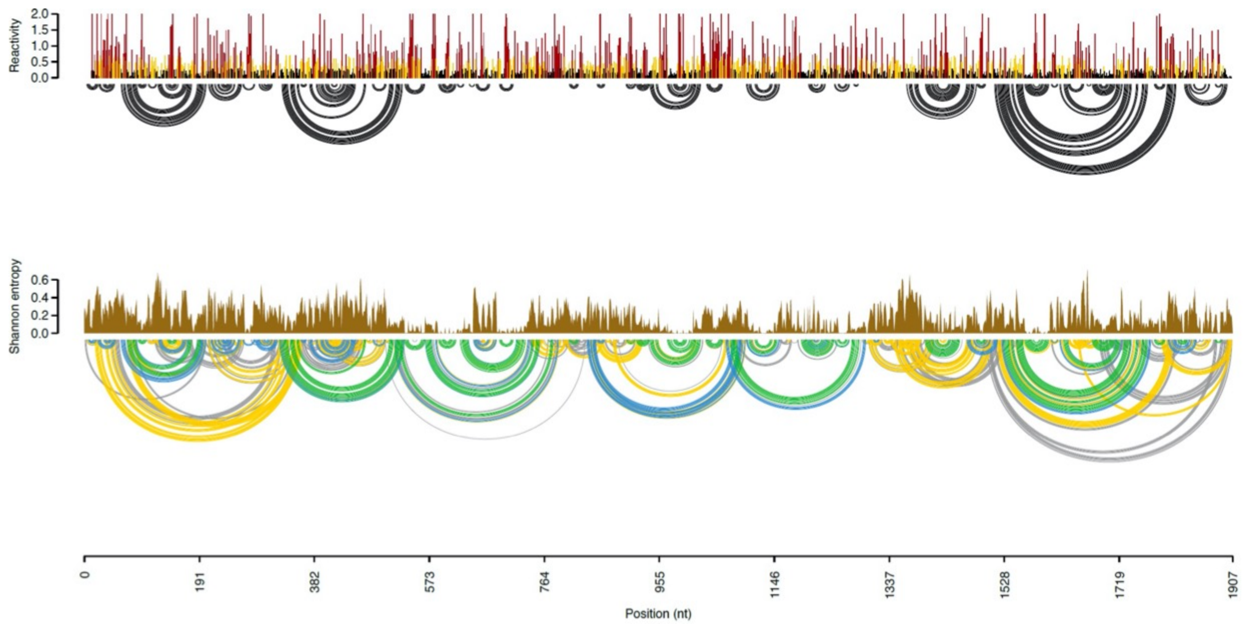


**Figure S8:** The SHAPE reactivity and Shannon entropy profiles for Chimeric luciferase reporter in the presence of compound **18.**

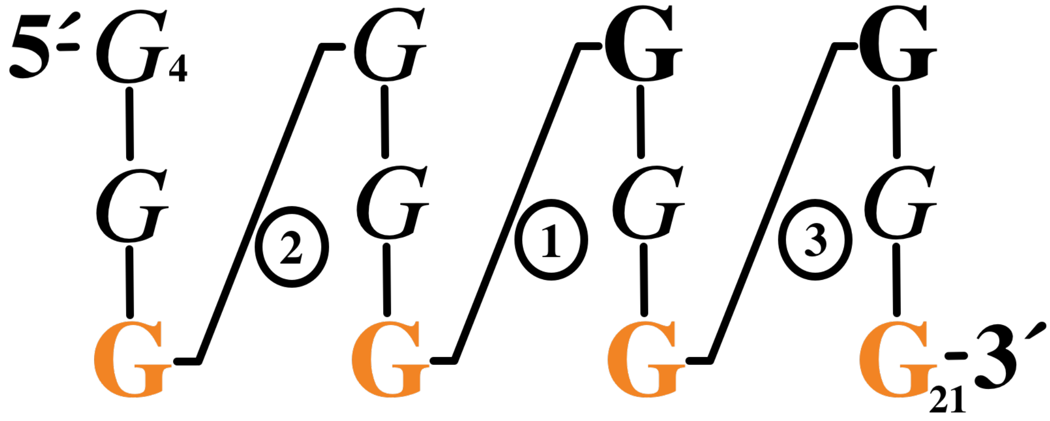


**Figure S9:** Schematic of the NRAS-G8U structure demonstrating the stereochemical properties adopted with in the G4 core. Description of the symbols used the schematic can be found in the review[13].

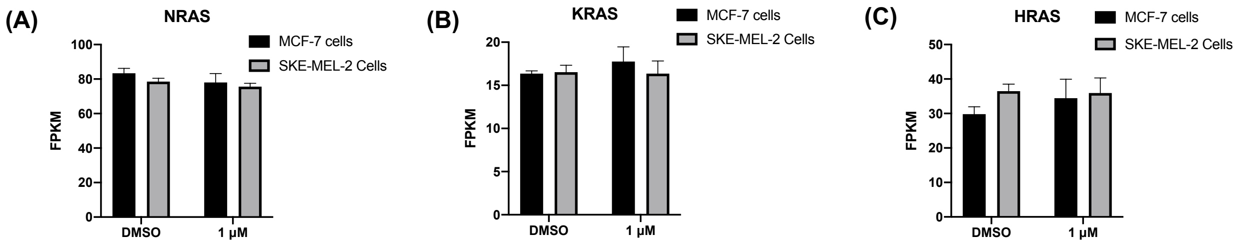


**Figure S10**: Bar graphs showing the FPKM values of other RAS family transcripts **(A)** NRAS **(B)** KRAS and **(C)** HRAS in the 18 treated and control samples.

(A)

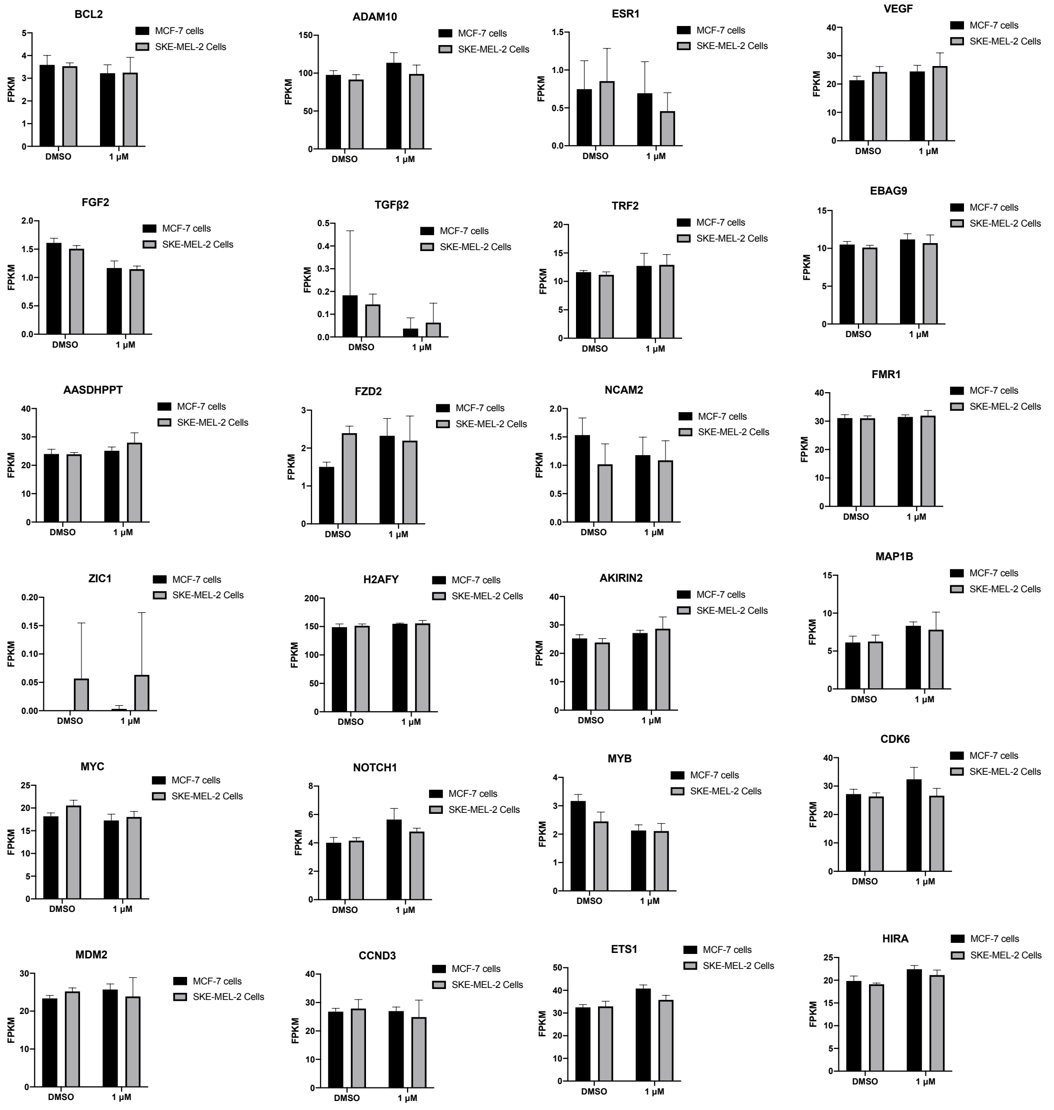


(B)

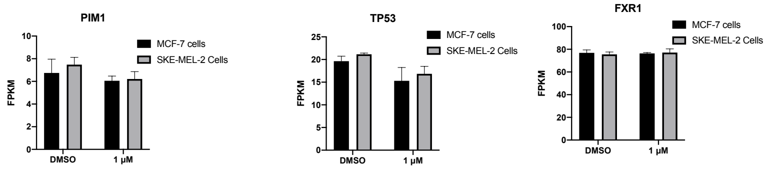


(C)

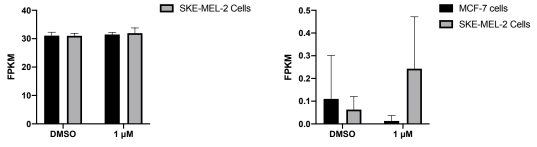


**Figure S11**: Bar graphs showing the FPKM values of other G4 containing transcripts in the 18 treated and control samples in the MCF-7 cells and SKE-MEL-2 cells **(A)**

Transcripts contain G4 at 5′UTR, **(B)** Transcripts contain G4 at 3′UTR **(C)** Transcripts contain G4 at ORF.

**Table S7:** Gene Ontology (GO) enrichment analysis of upregulated genes in MCF-7 cells (top) and SK-MEL-2 cells (bottom). GO term enrichment analysis based on Enrichr analysis of DESEQ2 reported genes with FDR < 0.05 and FC > 1.5. The top GO biological processes are listed along with their FDR adjusted p-value and odds ratio (OR).

| Index | Name | P-value | Adjusted p-value | Odds Ratio | Combined score |
|---|---|---|---|---|---|
| 1 | cellular heat acclimation (GO:0070370) | 0.000001990 | 0.0004703 | 218.71 | 2871.11 |
| 2 | heat acclimation (GO:0010286) | 0.000001990 | 0.0004703 | 218.71 | 2871.11 |
| 3 | intracellular sequestering of iron ion (GO:0006880) | 0.0002165 | 0.01050 | 144.22 | 1216.92 |
| 4 | positive regulation of nucleotide-binding oligomerization domain containing signaling pathway (GO:0070426) | 0.0002165 | 0.01050 | 144.22 | 1216.92 |
| 5 | regulation of hormone biosynthetic process (GO:0046885) | 0.0003238 | 0.01256 | 108.16 | 869.12 |
| 6 | regulation of nucleotide-binding oligomerization domain containing 2 signaling pathway (GO:0070432) | 0.0003238 | 0.01256 | 108.16 | 869.12 |
| 7 | positive regulation of extrinsic apoptotic signaling pathway via death domain receptors (GO:1902043) | 0.0003238 | 0.01256 | 108.16 | 869.12 |
| 8 | positive regulation of tau-protein kinase activity (GO:1902949) | 0.0003238 | 0.01256 | 108.16 | 869.12 |
| 9 | positive regulation of microtubule nucleation (GO:0090063) | 0.0004520 | 0.01619 | 86.53 | 666.41 |
| 10 | positive regulation of fibroblast migration (GO:0010763) | 0.0006008 | 0.01714 | 72.10 | 534.80 |

| Index | Name | P-value | Adjusted p-value | Odds Ratio | Combined score |
|---|---|---|---|---|---|
| 1 | intracellular sequestering of iron ion (GO:0006880) | 0.0001066 | 0.01472 | 207.61 | 1899.00 |
| 2 | positive regulation of extrinsic apoptotic signaling pathway via death domain receptors (GO:1902043) | 0.0001595 | 0.01472 | 155.70 | 1361.38 |
| 3 | L-serine biosynthetic process (GO:0006564) | 0.0001595 | 0.01472 | 155.70 | 1361.38 |
| 4 | regulation of hormone biosynthetic process (GO:0046885) | 0.0001595 | 0.01472 | 155.70 | 1361.38 |
| 5 | positive regulation of fibroblast migration (GO:0010763) | 0.0002965 | 0.02336 | 103.79 | 843.15 |
| 6 | positive regulation of coagulation (GO:0050820) | 0.0004745 | 0.02336 | 77.84 | 595.70 |
| 7 | positive regulation of hemostasis (GO:1900048) | 0.0004745 | 0.02336 | 77.84 | 595.70 |
| 8 | L-serine metabolic process (GO:0006563) | 0.0004745 | 0.02336 | 77.84 | 595.70 |
| 9 | erythrocyte development (GO:0048821) | 0.0005787 | 0.02336 | 69.18 | 515.75 |
| 10 | positive regulation of transcription from RNA polymerase II promoter in response to endoplasmic reticulum stress (GO:1990440) | 0.0005787 | 0.02336 | 69.18 | 515.75 |

**Table S8:** Gene Ontology (GO) enrichment analysis of down regulated genes in MCF-7 cells (top) and SK-MEL-2 cells (bottom) . GO term enrichment analysis based on Enrichr analysis of DESEQ2 reported genes with FDR < 0.05 and FC > -1.5. The top GO biological processes are listed along with their FDR adjusted p-value and odds ratio (OR).

| Index | Name | P-value | Adjusted p-value | Odds Ratio | Combined score |
|---|---|---|---|---|---|
| 1 | plasma membrane phospholipid scrambling (GO:0017121) | 0.005501 | 0.8201 | 22.89 | 119.07 |
| 2 | positive regulation of glycogen biosynthetic process (GO:0045725) | 0.01689 | 0.8201 | 11.44 | 46.69 |
| 3 | positive regulation of transmembrane transport (GO:0034764) | 0.01689 | 0.8201 | 11.44 | 46.69 |
| 4 | negative regulation of blood coagulation, intrinsic pathway (GO:2000267) | 0.07020 | 0.8201 | 17.11 | 45.44 |
| 5 | negative regulation of cell volume (GO:0045794) | 0.07020 | 0.8201 | 17.11 | 45.44 |
| 6 | fructose 2,6-bisphosphate metabolic process (GO:0006003) | 0.07020 | 0.8201 | 17.11 | 45.44 |
| 7 | fructose catabolic process (GO:0006001) | 0.07020 | 0.8201 | 17.11 | 45.44 |
| 8 | fructose catabolic process to hydroxyacetone phosphate and glyceraldehyde-3-phosphate (GO:0061624) | 0.07020 | 0.8201 | 17.11 | 45.44 |
| 9 | regulation of blood coagulation, intrinsic pathway (GO:2000266) | 0.07020 | 0.8201 | 17.11 | 45.44 |
| 10 | insulin metabolic process (GO:1901142) | 0.07020 | 0.8201 | 17.11 | 45.44 |

| Index | Name | P-value | Adjusted p-value | Odds Ratio | Combined score |
|---|---|---|---|---|---|
| 1 | plasma membrane phospholipid scrambling (GO:0017121) | 0.003403 | 0.7153 | 29.42 | 167.17 |
| 2 | SNARE complex assembly (GO:0035493) | 0.005389 | 0.7153 | 22.06 | 115.23 |
| 3 | positive regulation of glycogen biosynthetic process (GO:0045725) | 0.01058 | 0.7153 | 14.70 | 66.89 |
| 4 | negative regulation of blood coagulation, intrinsic pathway (GO:2000267) | 0.05524 | 0.7153 | 21.97 | 63.62 |
| 5 | negative regulation of cell volume (GO:0045794) | 0.05524 | 0.7153 | 21.97 | 63.62 |
| 6 | fructose 2,6-bisphosphate metabolic process (GO:0006003) | 0.05524 | 0.7153 | 21.97 | 63.62 |
| 7 | fructose catabolic process (GO:0006001) | 0.05524 | 0.7153 | 21.97 | 63.62 |
| 8 | fructose catabolic process to hydroxyacetone phosphate and glyceraldehyde-3-phosphate (GO:0061624) | 0.05524 | 0.7153 | 21.97 | 63.62 |
| 9 | regulation of blood coagulation, intrinsic pathway (GO:2000266) | 0.05524 | 0.7153 | 21.97 | 63.62 |
| 10 | insulin metabolic process (GO:1901142) | 0.05524 | 0.7153 | 21.97 | 63.62 |

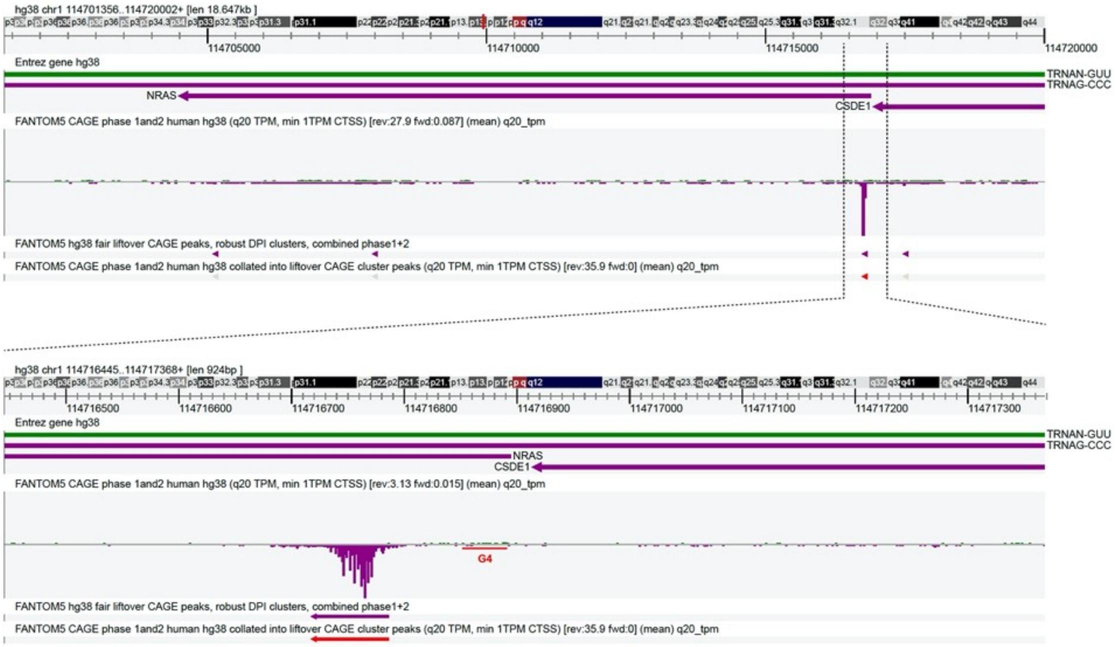

**Figure S12:** ZENBU analysis of the NRAS TSS. The G4 regions marked with red underline and are in the promoter region.

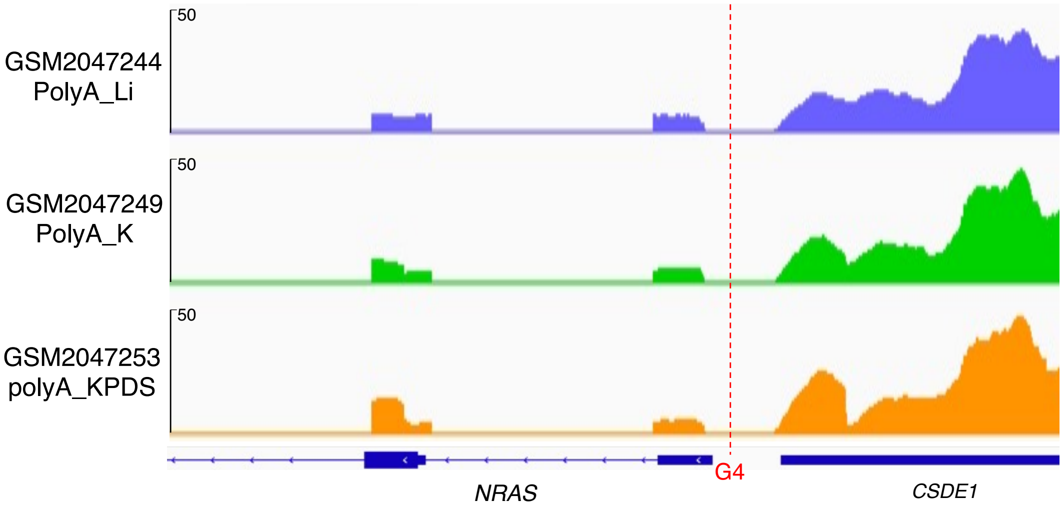


**Figure S13:** The probability of NRAS G4 formation in the 5' UTR of NRAS mRNA obtained by genome-wide rG4-seq analysis; the highlighted region represents NRAS G4 that is present in the promoter region of the NRAS gene but not in the 5' UTR.

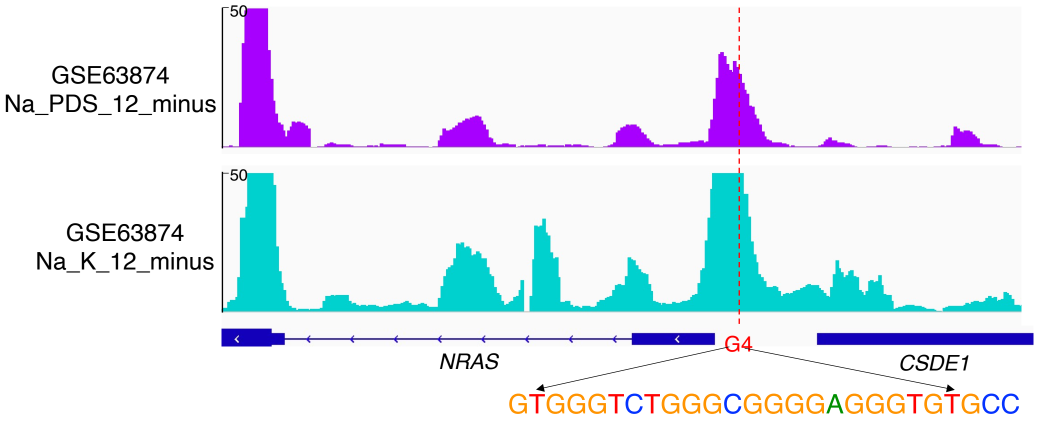


**Figure S14**: The probability of NRAS G4 formation in the promoter region of NRAS gene obtained by genome-wide G4-seq analysis; the highlighted region represents NRAS G4 that is presence in the promoter region of NRAS gene.

**Table S9**: The sequence information of the primers used in the RT-PCR.

| Name | Sequence (5′ to 3′) |
|---|---|
| A | AAC GTC CCG TGT GGG AGG GG |
| B | AGA CCC CGG AAC CGC CAT GA |
| C | CCT ATA CAA TGT ATG TAA TTT GTT TCC |
| D | CAA TGC ACC AAA GTT TTA CAA TAT TTG AAC |
| E | TGTGGATTTCGAGTCGTCTTAAT |
| F | CGAAGAAGGAGAATAGGGTTGG |
